# Engineered High-Affinity ACE2 Peptide Mitigates ARDS and Death Induced by Multiple SARS-CoV-2 Variants

**DOI:** 10.1101/2021.12.21.473668

**Authors:** Lianghui Zhang, Soumajit Dutta, Shiqin Xiong, Matthew Chan, Kui K. Chan, Timothy M. Fan, Keith L. Bailey, Matthew Lindeblad, Laura M. Cooper, Lijun Rong, Anthony F. Gugliuzza, Diwakar Shukla, Erik Procko, Jalees Rehman, Asrar B. Malik

**Author notes:** Corresponding authors: Asrar B. Malik, Jalees Rehman, Erik Procko, Diwakar Shukla, or Lianghui Zhang. These authors contributed equally (Soumajit Dutta, Shiqin Xiong, Matthew Chan).

## Abstract

Vaccine hesitancy and continuing emergence of SARS-CoV-2 variants of concern that may escape vaccine-induced immune responses highlight the urgent need for effective COVID-19 therapeutics. Monoclonal antibodies used in the clinic have varying efficacies against distinct SARS-CoV-2 variants; thus, there is considerable interest in engineered ACE2 peptides with augmented binding affinities for SARS-CoV-2 Spike protein. These could have therapeutic benefit against multiple viral variants. Using molecular dynamics simulations, we show how three amino acid substitutions in an engineered soluble ACE2 peptide (sACE2_2_.v2.4-IgG1) markedly increase affinity for the SARS-CoV-2 Spike (S) protein. We demonstrate high binding affinity to S protein of the early SARS-CoV-2 WA-1/2020 isolate and also to multiple variants of concern: B.1.1.7 (Alpha), B.1.351 (Beta), P.1 (Gamma), and B.1.617.2 (Delta) SARS-CoV-2 variants. In humanized K18-hACE2 mice, prophylactic and therapeutic administration of sACE2_2_.v2.4-IgG1 peptide prevented acute lung vascular endothelial injury and lung edema (essential features of ARDS) and significantly improved survival after infection by SARS-CoV-2 WA-1/2020 as well as P.1 variant of concern. These studies demonstrate for the first time broad efficacy *in vivo* of an ACE2 decoy peptide against multiple SARS-CoV-2 variants and point to its therapeutic potential.

## INTRODUCTION

The COVID-19 pandemic, caused by severe acute respiratory syndrome-associated coronavirus 2 (SARS-CoV-2), has resulted in over 3.5 million deaths world-wide by June 2021. The successful development of mRNA-based and adenoviral vaccines, which induce immunity against the SARS-CoV-2 Spike (S) protein with reported efficacy rates of 70% to 95% ^1–7^, represent a monumental milestone in reducing the pandemic. However, persistence of vaccine hesitancy in high-income countries ^8^, limited supply of vaccines in low-income countries, breakthrough infections, and emergence of B.1.1.7 (“Alpha”), B.1.351 (“Beta”), P.1 (“Gamma”), and B.1.617.2 (“Delta”) SARS-CoV-2 variants of concern (VOC), which carry S protein mutations exhibiting higher affinity for ACE2 that may escape vaccine-induced humoral immune responses ^9–11^, highlights the urgent need for novel targeted therapeutics.

The trimeric S protein on the SARS-CoV-2 surface is a class I fusion protein proteolytically processed during biosynthesis into soluble S1 and membrane-tethered S2 subunits that remain non-covalently associated ^12–14^. The S1 subunit contains the receptor-binding domain (RBD), which engages ACE2 as a cell-surface entry receptor ^12, 14, 15^. The critical role of S protein for viral entry has led to the search for anti-SARS-CoV-2 therapeutics targeting the S-ACE2 interaction, such as monoclonal antibodies (mAbs) binding to epitopes on S that neutralize virus infection ^16, 17^.

Monoclonal antibodies not only provide tight affinity for the viral target (K_D_ < 1 nM) but also harness endogenous host defense mechanisms, including interactions with the neonatal Fc receptor (FcRn) to confer long serum stability (typical half-lives of IgG1 proteins are several weeks) and with FcγR on immune cells to facilitate SARS-CoV-2 clearance in vivo ^18^. However, there are also serious limitations of mAb therapies. First, emergence of VOCs with S protein mutations appears to diminish neutralization capacity in some cases ^19, 20^. Second, mAbs are typically administered by intravenous (IV) infusion and less is known about their pharmacokinetic stability when applied locally in lungs via inhalation.

An alternative to mAbs is to use soluble wild type ACE2 (sACE2) or engineered soluble ACE2 peptides (we refer here to soluble constructs of ACE2 that include both protease and dimerization domains, sACE2_2_), which function as decoys for the viral S protein^21, 22^ . Wild type sACE2_2_ has shown indications of efficacy in the clinic ^23^ but was not effective as an Fc fusion for neutralizing virus in an *in vivo* infection model ^24^, and there are opportunities for next generation ACE2 derivatives with improved properties. We used deep mutagenesis to show that many amino acid substitutions within ACE2 enhanced the affinity for S protein ^22^. We also showed that a derivative of sACE2_2_ with 3 mutations (T27Y, L79T, and N330Y), called sACE2_2_.v2.4, bound S with 35-fold tighter affinity (K_D_ = 0.6 nM) and neutralized authentic SARS-CoV-2 with sub-nanomolar IC50, on par with high affinity mAbs ^22^. We showed that affinity increases (K_D_ < 0.1 nM) for S variants carrying the N501Y mutation associated with increased virus transmission ^25^. However, studies have not examined the efficacy of sACE2_2_.v2.4 *in vivo* and its capacity to bind S protein from the Alpha, Beta, Gamma, and Delta VOCs.

Laboratory mouse strains do not show significant disease pathology when infected with SARS-CoV-2 because murine ACE2 has low affinity for SARS-CoV-2 S protein ^26^. The K18-hACE2 transgenic mouse model expressing human ACE2 driven by the epithelial K18 promoter developed severe lung injury and immune dysregulation following SARS-CoV-2 inoculation ^27^. Acute lung injury (ALI) or its progression to severe acute respiratory distress syndrome (ARDS) observed in critically ill COVID-19 patients is characterized by endothelial injury and breakdown of lung vascular barrier function via loss of lung endothelial cells and disruption of endothelial adherens junction protein VE-cadherin, which lead to protein-rich lung edema and respiratory failure ^28–30^. However, it is not clear whether animal models of COVID-19 show an analogous pathology of the lung vascular endothelium and whether treatment following inoculation with SARS-CoV-2 variants can prevent lung endothelial injury and ARDS or reduce mortality.

Here, we found that inoculation of K18-hACE2 mice with an original SARS-CoV-2 variant (WA isolate) induced severe lung vascular injury and that inoculation with the P.1 variant induced even greater injury as well as higher mortality. We explored the molecular basis for the tight binding between S of SARS-CoV-2 and our sACE2_2_.v2.4 decoy peptide by molecular dynamics (MD). We also characterized pharmacokinetics (PK) of sACE2_2_.v2.4-IgG1 in mice via different routes and further showed that tight affinity of sACE2_2_.v2.4 persisted against newly emerging SARS-CoV-2 VOCs. sACE2_2_.v2.4-IgG1 showed exceptional efficacy in reducing viral entry, lung vascular hyperpermeability and preventing ARDS and mortality on challenging humanized mice with SARS-CoV-2 WA-1/2020 or the P.1 VOC.

## RESULTS

### Comparison of acute lung endothelial injury and ARDS induced by SARS-CoV-2 WA-1/2020 and P.1 variant of concern in K18-hACE2 transgenic mice

We inoculated K18-hACE2 mice intranasally with live SARS-CoV-2 WA-1/2020 virus at 1×10^4^ PFU (N=10) or 1×10^5^ PFU (N=10), and observed body weight loss, a marker of severe viral infection and survival. The higher dose (1×10^5^ PFU) induced mortality starting at Day 5 whereas mice in the lower dose group (1×10^4^ PFU) reached humane endpoints or died starting on Day 7 (**Figure 1A**). The degree of weight loss was comparable with both viral doses during the first 5 days but due to the rapid death in the high dose group, we were unable to compare weight losses in both groups during advanced stages of infection (**Figure 1B**). We therefore chose inoculation with the lower dose (1×10^4^ PFU) of SARS-CoV-2 WA-1/2020 to study lung vascular injury as the disease progressed to the advanced stage. Lung transvascular permeability was evaluated at Day 7 post-inoculation of SARS-CoV-2 isolate WA-1/2020 (1×10^4^ PFU) using intravenous (IV) infusion of Evans Blue Albumin (EBA) tracer. Representative lung images (**Figure 1C**) and quantification of lung EBA (lung microvessel permeability of albumin) (**Figure 1D**) showed that SARS-CoV-2 induced severe lung vascular endothelial injury at Day 7. Quantification of lung wet/dry ratio (a measure of pulmonary edema) indicated that SARS-CoV-2 also induced severe lung edema by Day 7 (**Figure 1E**). Flow cytometric quantification of lung endothelial cells (ECs) showed that one third of the lung EC population was lost by Day 7 (**Figure 1F-G**). Furthermore, quantitative real-time PCR showed that the mRNA levels of the endothelial adherens junction protein VE-cadherin significantly decreased by Day 7(**Figure 1H**). Histological analysis of lungs showed pulmonary hemorrhage and edema at Day 7 (**Figure 1I**) and massive infiltration of immune cells (**Figure 1I**). Fibrosis staining (Trichrome blue and Sirius red) showed that SARS-CoV-2 infection resulted in collagen deposition at Day 7 (**Figure 1I**).

**Figure 1.**
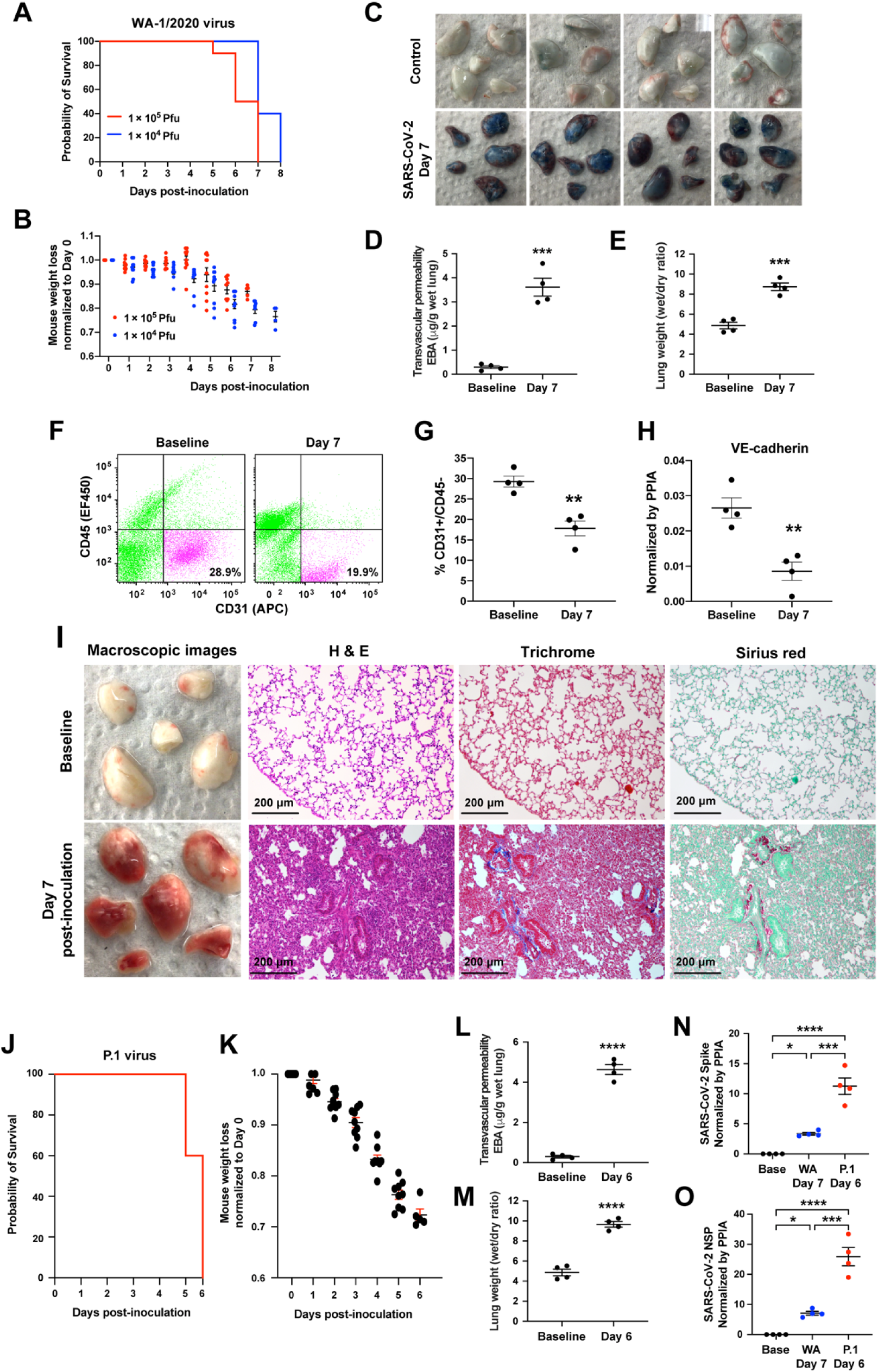
SARS-CoV-2 isolates WA-1/2020 and P.1 (Brazil) induce severe lung vascular endothelial injury and ARDS in K18-hACE2 transgenic mice. (**A-B**) K18-hACE2 transgenic mice were inoculated with SARS-CoV-2 isolate WA-1/2020 at 1×10^4^ PFU and 1×10^5^ PFU respectively. N=10. The mice were observed for the survival (**A**) and weight (**B**). (**C-J**) The K18-hACE2 transgenic mice were inoculated with SARS-CoV-2 WA-1/2020 at 1×10^4^ PFU. The mice were harvested at Day 7 using non-infected baseline mice as controls. N=4. (**C**) Macroscopic images of lungs following Evans Blue Albumin (EBA) injection with subsequent perfusion to remove EBA in circulation. Quantification of EBA as a marker of pulmonary transendothelial permeability (**D**) and of lung wet/dry ratio as a measure of lung edema (**E**). The representative plots (**F**) and flow cytometry analysis (**G**) of CD31+/CD45- lung ECs as a percentage of total lung cells at baseline and day 7 post-inoculation. (**H**) Quantification of Cdh5 mRNA expression normalized by the housekeeping gene PPIA (peptidylpropyl isomerase A). (**I**) Macroscopic lung images (1^st^ column), histology images of lung sections with H & E staining (2^nd^ column), and Masson Trichrome (3^rd^ column) and Sirius Red (4^th^ column) staining for collagen deposition at baseline (1^st^ row) and day 7 post-inoculation (2^nd^ row). (**J-O**) K18-hACE2 transgenic mice were inoculated with SARS-CoV-2 P.1 variant at 1×10^4^ PFU. N=10. The mice were observed to assess survival (**J**) and weight loss (**K**). The mice were harvested at Day 6 to evaluate lung vascular leak by EBA assay (**L**) and lung wet/dry ratio (**M**). N=4. (**N**) Real-time quantitative PCR was applied to measure viral loads in the lungs of SARS-CoV-2 P.1 variant at Day 6 post-inoculation and the WA-1/2020 strain at Day 7 post-inoculation. The mRNA expression levels of SARS-CoV-2 Spike (**N**) and SARS-CoV-2 NSP (**O**) are shown. Data are presented as mean ± SEM. **: P<0.01, ***: P<0.001, ****: P<0.0001 by Student’s t-test for **D, E, G, H, L**, and **M**; One-way ANOVA for **N** and **O**.

We also inoculated K18-hACE2 mice with the P.1 variant at a dose of 1×10^4^ PFU and observed death beginning earlier at Day 5 (**Figure 1J**). The P.1 variant induced significantly greater weight loss (**Figure 1K & 1B**), lung vascular hyper-permeability (**Figure 1L**) and edema (**Figure 1M**) by Day 6. We assessed viral replication by quantifying the expression level of the SARS-CoV-2 Spike protein and SARS-CoV-2 non-structural protein (NSP) and found that detectable levels of P.1 variant at Day 6 post-inoculation were 5 times greater than of WA-1/2020 at Day 7 (**Figure 1N-O**). Thus, SARS-CoV-2 isolate WA-1/2020 and P.1 VOC induced severe acute lung endothelial injury, resulting in lung transvascular permeability, lung edema, and death. P.1 variant induced more severe lung vascular injury and ARDS and death occurred 2 days earlier as compared to WA-1/2020.

### Molecular basis for tight binding of sACE2_2_.v2.4 to SARS-CoV-2 Spike protein

We next focused on a high affinity soluble ACE2 peptide that had been engineered to therapeutically target the S protein across multiple SARS-CoV-2 variants. Assessment of the relative binding activities of single amino acid substitutions has previously informed design of the high-affinity peptide sACE2_2_.v2.4 ^22^. To better understand the molecular basis for the increased affinity to the viral Spike protein, wild type and v2.4 ACE2 proteins were simulated by all-atom MD in both the unbound/apo and RBD-bound states. Approximately 400 µs of simulation data were collected for each apo protein. For all residues within the interface, the backbone and sidechain distances were calculated with respect to all other residues within 0.8 nm. A total of 74 distances were computed, and to assess surface dynamics, time-lagged independent component analysis (TICA) determined critical residue movements from the higher dimensional dataset. TICA linearly transformed the MD dataset to find the slowest motions in the surface. For both wild type and v2.4 ACE2, three TICs were found with timescales larger than the lag time (**Figures S1A-C**). The distance features that showed the highest correlation with the three TICs are the same for both proteins (**Figures S1A-B**) and are near glycosylation sites (ACE2 residues 103 and 546). The projection of TIC 1 and TIC 2 components showed that these motions have similar distributions (**Figures S1D-E**), suggesting that residue dynamics at the surface are not substantially changed between the two proteins.

To examine how the v2.4 mutations impacted interfacial interactions, we performed ∼530 µs of simulation for wild type and v2.4 ACE2 proteins with the RBD bound. TICA calculations on interface residue distances ^35^ showed that features most correlated with the four slowest components are in RBD loops 1 (residues 477-487) and 2 (residues 497-505) (**Figure S2**). RBD loop 1 is close to v2.4 mutations T27Y and L79T, while RBD loop 2 is close to v2.4 mutation N330Y. Trajectory observation revealed that new polar interactions formed in the RBD loop regions with ACE2.v2.4 that were not present in the wild type complex. Y27 of ACE2.v2.4 hydrogen bonds to Y473 in RBD loop 1, ACE2.v2.4-T79 interacted with the backbone carbonyl of RBD-G485, and ACE2.v2.4-Y330 interacted with the backbone carbonyl of RBD-P499 (**Figure 2A**). To obtain the ensemble distance distributions of these stable interactions, we built a Markov state model (MSM) on the MD data. MSM-weighted distributions demonstrated that all three newly formed interactions in the ACE2.v2.4 complex were stable (distance distributions are single peaks within 0.6 nm; **Figure 2B**) and that the RBD loops are stabilized in closer positions in ACE2.v2.4 compared to the wild type complex (**Figures 2C-D**). Furthermore, angle calculations for the loops with respect to the centers of mass for the RBD and ACE2 proteins demonstrated reduced variance in ACE2.v2.4 (RBD loop 1 standard deviation [SD] 2.0°, loop 2 SD 1.0°) compared to wild type (loop 1 SD 2.9°, loop 2 SD 1.9°). Overall, new interactions formed by ACE2.v2.4 stabilized RBD loops in close association with the receptor surface and higher affinity was explained by localized effects at the interface.

**Figure 2.**
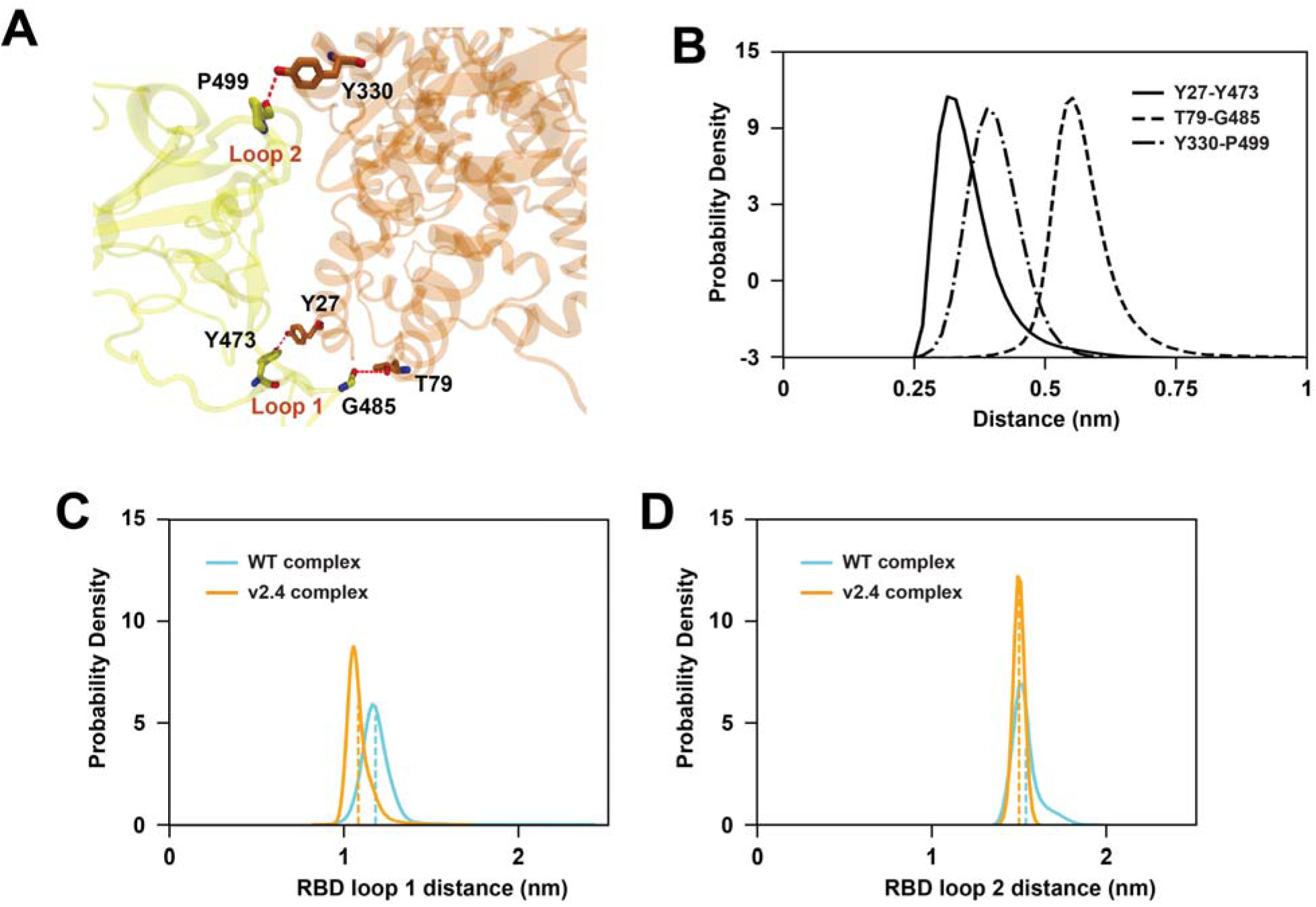
Mutations in ACE2_2_.v2.4 form new interactions stabilizing RBD loops. (**A**) RBD-bound ACE2 proteins were simulated. Newly formed polar interactions between ACE2.v2.4 (orange) and RBD (yellow) are indicated by broken red lines. (**B**) MSM-weighted distance distributions of newly formed polar interactions between RBD and ACE2.v2.4. (**C**-**D**) Distance distributions between the centers of mass of RBD loops 1 and 2 with respect to ACE2 residues (Cα atoms) 27 (**C**) and 330 (**D**), respectively. Loop centers of mass are calculated from Cα atoms. The means of each distribution are shown by vertical dashed lines. Distributions from simulations of RBD-bound wild type ACE2 are blue, RBD-bound ACE2.v2.4 are orange.

### Pharmacokinetics of sACE2_2_.v2.4-IgG1 peptide

Prior to performing *in vivo* efficacy studies using the sACE2_2_.v2.4-IgG1 peptide described below, we characterized the pharmacokinetics (PK) of the peptide in mice. We injected sACE2_2_.v2.4 (0.5 mg/kg) without a fusion partner into the tail veins of male and female mice and found the protein was rapidly cleared with a serum half-life estimated to be under 10 minutes as measured by ACE2 ELISA (**Figure S3A**) and ACE2 catalytic activity in serum (**Figure S3B**). This is much shorter than the serum half-life of wild-type sACE2_2_ in humans (of 2 to 3 hours) ^36^. No toxicity was observed when sACE2_2_.v2.4 was IV administered twice daily at 0.5 mg/kg for five consecutive days. Mice were euthanized on Day 7 and blood chemistry, hematology, and tissue pathology showed no differences with mock treated mice, underscoring the safety of sACE2_2_.v2.4.

Using an ELISA for human IgG1, both wild type sACE2_2_-IgG1 and sACE2_2_.v2.4-IgG1 showed equivalent serum PK after IV administration (2.0 mg/kg) in male mice, with protein persisting for over 7 days (**Figure S3C**). We therefore concluded that the three mutations in the high affinity sACE2_2_.v2.4 derivative did not substantially change PK. We next examined PK in male and female mice and again saw human IgG1 protein persisting for days in the serum (**Figure 3A**), yet the ACE2 moiety was cleared within 24 hours based on an ACE2 ELISA (**Figure 3B**). Measurement of ACE2 catalytic activity revealed even faster decay (**Figure 3C**). Immunoblot for human IgG1 confirmed that the fusion protein was proteolyzed in mouse blood to liberate long-lived IgG1 fragments (**Figure 3D**). Clearance of sACE2_2_.v2.4-IgG1 was recapitulated *in vitro* by incubating the protein with normal mouse serum over days (**Figure 3E**), but the protein was resistant to degradation in human serum (**Figure 3F**), suggesting that human sACE2_2_ has better stability in human patients and mouse studies likely underestimated how the protein will perform in humans. Further, no toxicity was observed in mice IV administered sACE2_2_.v2.4-IgG1 (2.0 mg/kg), followed up 7 days later with blood chemistry, hematology, and tissue pathology analysis.

**Figure 3.**
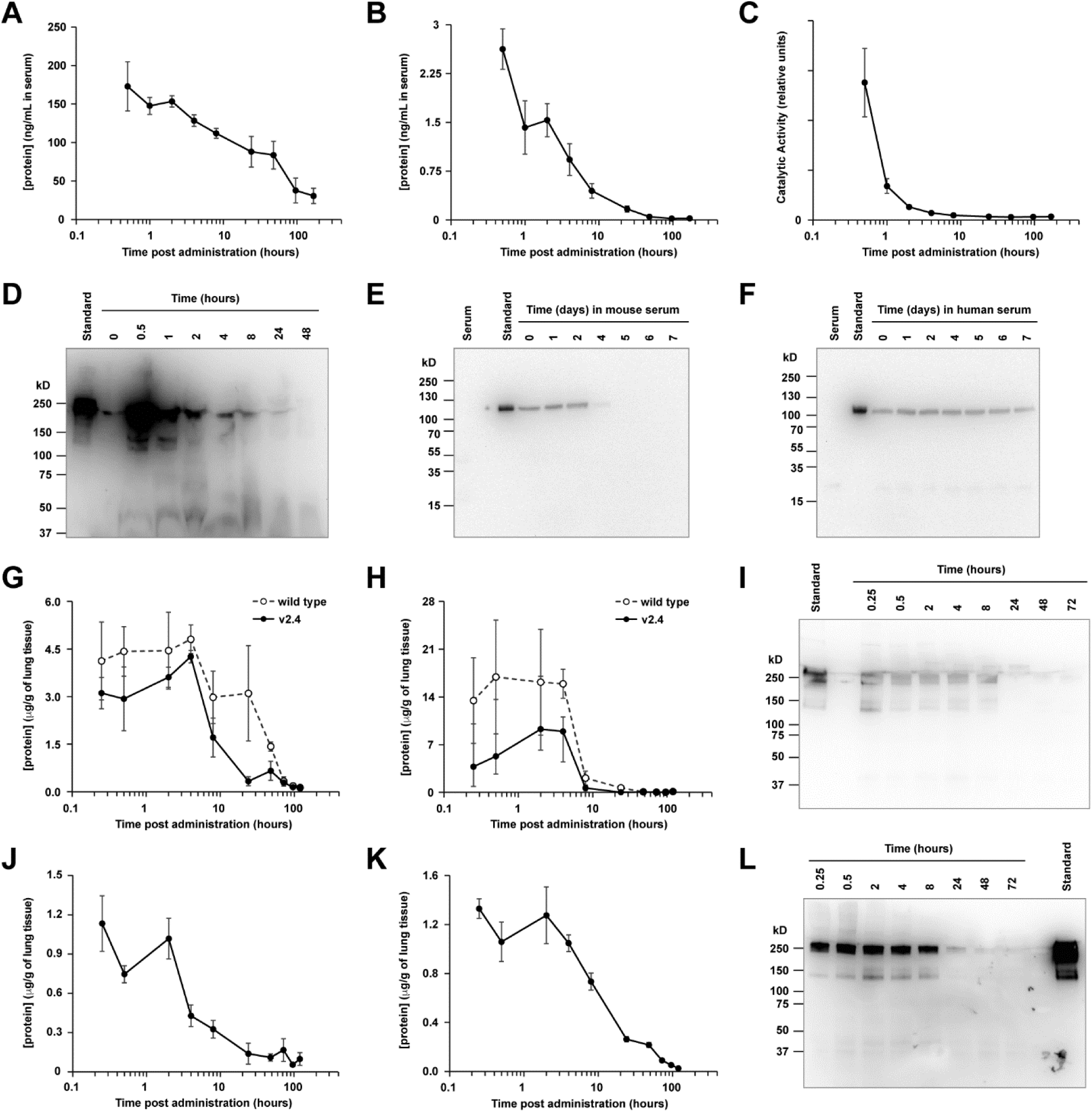
Pharmacokinetics of sACE2_2_.v2.4-IgG1 peptide. (**A**-**C**) sACE2_2_.v2.4-IgG1 was IV administered to mice (N=6 per time point; 2.0 mg/kg). Serum was collected and analyzed by human IgG1 ELISA (**A**), by ACE2 ELISA (**B**), and for ACE2 catalytic activity (**C**). (**D**) Serum samples from representative male mice were separated on a non-reducing SDS electrophoretic gel and probed with anti-human IgG1 using 10 ng of purified sACE2_2_.v2.4-IgG1 as a standard. Predicted molecular weight (MW; excluding glycans) of dimer is 216 kD. (**E**-**F**) sACE2_2_.v2.4- IgG1 was incubated in vitro at 37 °C with normal mouse (**E**) and human (**F**) serum. Samples at the indicated time points were separated on a reducing SDS gel and immunoblotted with anti-human ACE2. MW of monomer (excluding glycans) is 108 kD. Shown are representative blots from two experiments. (**G**-**H**) Wild type sACE2_2_-IgG1 (white circles) and sACE2_2_.v2.4-IgG1 (black circles) were administered IT at 1.0 mg/kg. Lung tissues were collected, and proteins were extracted and analyzed by (**G**) human IgG1 ELISA and (**H**) ACE2 ELISA. N=3 males per time point. (**I**) Lung extracts from representative mice IT administered sACE2_2_.v2.4-IgG1 were analyzed under non-reducing conditions by anti-human IgG1 immunoblot. (**J**-**K**) Mice inhaled nebulized sACE2_2_.v2.4-IgG1. Extracts from lung tissue were analyzed by (**J**) ACE2 ELISA and (**K**) human IgG1 ELISA. N=3 males per time point. (**L**) Representative extracts from lung tissue of mice receiving nebulized sACE2_2_.v2.4-IgG1 were analyzed by anti-human IgG1 immunoblot. Data are presented as mean ± SEM.

In addition, the effects of sACE2_2_.v2.4-IgG1 delivered directly to the respiratory tract, the primary site of SARS-CoV-2 infection, was studied. Following intratracheal (IT) delivery (1.0 mg/kg), sACE2_2_.v2.4-IgG1 persisted in lungs for at least 4 hours determined by ACE2 ELISA, human IgG1 ELISA, and anti-human IgG1 immunoblot (**Figure 3G-I**). Levels of sACE2_2_.v2.4-IgG1 absorbed into the blood were too low for detection. As observed for IV administration, wild type sACE2_2_-IgG1 and sACE2_2_.v2.4-IgG1 had similar PK in lungs following IT delivery. We further investigated administration of sACE2_2_.v2.4-IgG1 by inhalation, in which the protein was nebulized for 30 minutes into a chamber holding the mice. sACE2_2_.v2.4-IgG1 remained high and relatively constant for 2 hours (**Figure 3J-L**). The different PK profiles based on route of administration (i.e., protein delivered directly into lungs persisted for hours but did not reach detectable levels in plasma, whereas IV delivered protein achieved high but short-lived plasma concentrations) offer opportunities to treat SARS-CoV-2 in different ways, guided by whether SARS-CoV-2 is primarily located in the respiratory tract or disseminated systemically.

### Engineered sACE2_2_.v2.4-IgG1 prevents SARS-CoV-2 entry and SARS-CoV-2-induced acute lung injury

We next assessed whether sACE2_2_.v2.4-IgG1 blocked entry of SARS-CoV-2 into cells using a replication-deficient pseudovirus which expresses the S protein and thus models SARS-CoV-2 cell entry ^19^. Entry is mediated by S engagement of the surface receptor ACE2 and the surface protease TMPRSS2, as well as other entry mediators that facilitate infection ^13, 37, 38^. We studied three cell types: the human lung epithelial A549 cell line, primary human lung microvascular endothelial cells (hLMVECs), and an A549 cell line stably over-expressing hACE2 (hACE2-A549) to maximize its infectability by the pseudovirus. We observed that ACE2 and TMPRSS2 were expressed in all three cell types albeit at significantly higher levels in hACE2-A549 cells and lower levels in hLMVECs (**Figure 4A**). After pre-incubation of each cell type with sACE2_2_-IgG1 (i.e., wild-type) or sACE2_2_.v2.4-IgG1 at 5 or 25 µg/ml for 1 hour, luciferase-expressing pseudovirus was added (MOI=0.1) for 24 hours. Luciferase activity in the cells after removal of the replication-deficient pseudovirus served as a measure of viral entry. Pre-treatment with sACE2_2_-IgG1 significantly reduced SARS-CoV-2 pseudovirus entry in all three cell types but pre-treatment with sACE2_2_.v2.4-IgG1 showed far greater efficacy in blocking viral entry (**Figure 4B**). sACE2_2_.v2.4-IgG1 at 5 µg/ml was as effective as 25 µg/ml of sACE2_2_-IgG1, suggesting that the engineered peptide had 5-fold greater efficacy in this assay.

**Figure 4.**
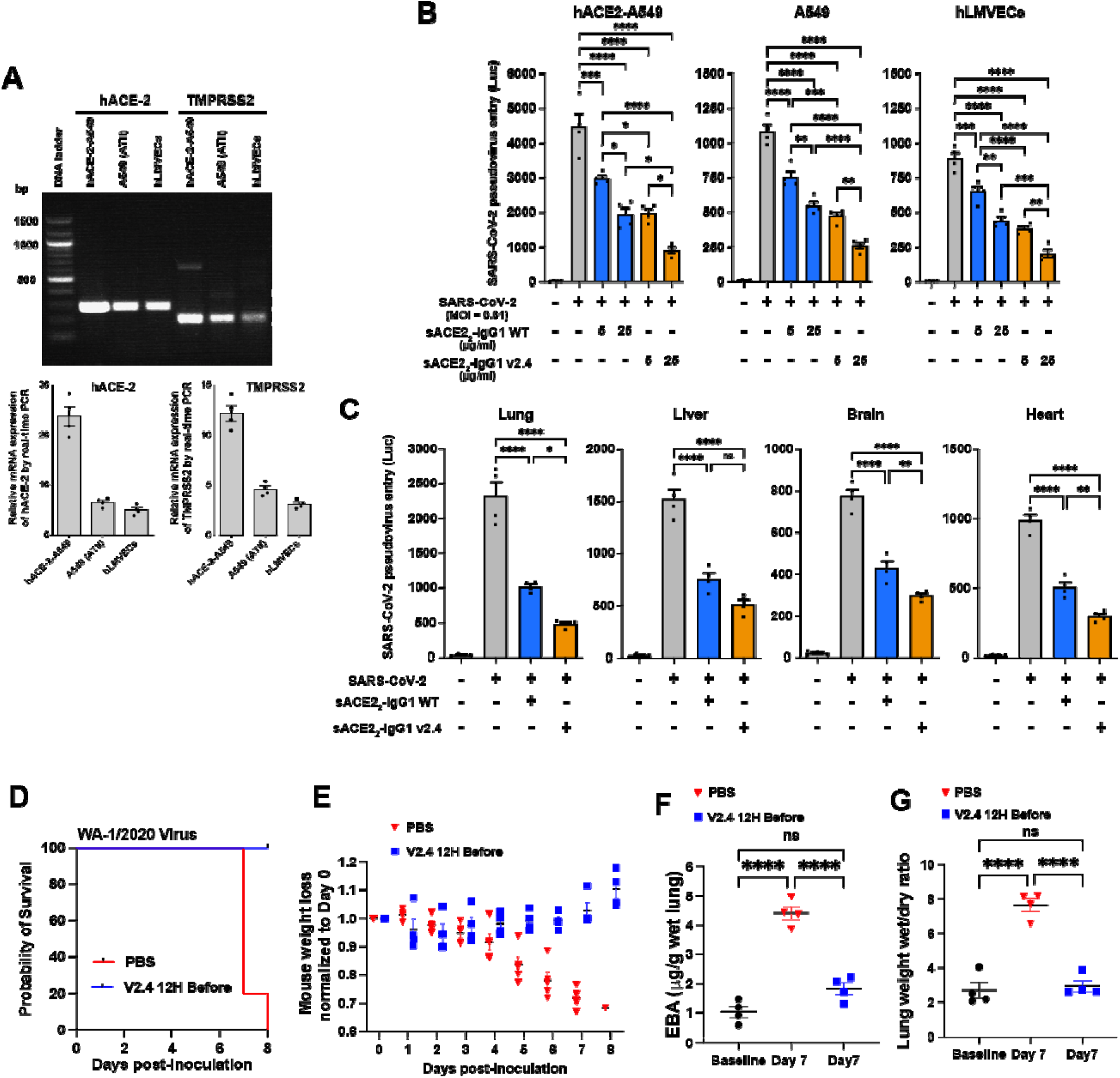
Engineered sACE2_2_.v2.4-IgG1 peptide prevents SARS-CoV-2 pseudovirus cell entry and SARS-CoV-2-induced acute lung injury. (A) mRNA expression levels of ACE-2 (NM_001371415.1) and TMPRSS2 (NM_001135099) in human lung epithelial A549 cells, A549 cells that stably express hACE2and hLMVECs (human lung microvascular endothelial cells) were analyzed by one-step RT-PCR (above) and real-time PCR (below). Relative expression was normalized to GAPDH expression levels. **(B)** Cultured hACE2-A549, A549, and hLMVECs were preincubated with sACE2_2_-IgG1 or sACE2_2_.v2.4-IgG1 at 5 or 25 µg/ml for 1 hour. SARS-CoV-2 pseudovirus (MOI=0.1) was added to the cells and the cells were harvested at 24 hours. Virus entry was evaluated by luciferase activity. N = 4 replicates. **(C)** 10 mg/kg sACE2_2_-IgG1, sACE2_2_.v2.4-IgG1, or peptide buffer (PBS + 0.2% BSA) was intravenously administrated into K-18 hACE2 transgenic mice for 30 minutes prior to SARS-CoV-2 pseudo-entry virus (10^6^ pfu) i.p. injection. Tissue lysates were prepared at 24h and virus entry in the selected organs was evaluated by luciferase activity. Peptide buffer was applied as control group. N=4. (**D** - **G**) K18-hACE2 transgenic mice were inoculated with SARS-CoV-2 isolate WA-1/2020 at 1×10^4^ PFU. The mice received control PBS or sACE2_2_.v2.4-IgG1 10mg/kg via IV injection 12h before inoculation. Mice were observed for survival (**D**) and body weight (**E**), N=5. Quantification of EBA as a marker of pulmonary transendothelial permeability (**F**) and lung wet/dry ratio (**G**) as a measure of lung edema. Data are mean ± SEM, N=4. *P<0.05, **: P<0.01, ***: P<0.001, ****: P<0.0001 by one-way ANOVA. ns: not significant.

We next examined *in vivo* efficacy. Based on the PK studies above, 10 mg/kg sACE2_2_-IgG1 (WT), 10 mg/kg sACE2_2_.v2.4-IgG1, or control peptide buffer (PBS + 0.2% bovine serum albumin [BSA]) was injected IV into K18-hACE2 transgenic mice at 30 minutes prior to systemic pseudovirus (10^6^ PFU i.p.) administration and mice were then assessed at 24 hours. Tissue lysates of lung, heart, brain, and liver (**Figure 4C**) were assessed for luciferase activity. Viral entry into lungs was substantially higher than other organs even though the pseudovirus was injected intraperitoneally, consistent with preferred viral entry into type II alveolar lung epithelium due to higher hACE2 expression^39^. Inhibition of pseudovirus entry was significantly greater in sACE2_2_.v2.4-IgG1 pre-treatment mice when compared to sACE2_2_-IgG1 (WT) pre-treatment group (**Figure 4C)**.

To test the prophylactic effects of sACE2_2_.v2.4-IgG1 on live SARS-CoV-2 infection, K18-hACE2 transgenic mice were pretreated with a single dose of sACE2_2_.v2.4-IgG1 (10mg/kg) via IV injection 12h before SARS-CoV-2 isolate 1/2020 WA (1×10^4^ PFU) inoculation. While the PK studies indicated that serum levels of sACE2_2_.v2.4-IgG1 in mice fall within hours, investigations of ‘long-lived’ monoclonal antibodies with similar affinities for the viral spike have demonstrated prophylaxis with protein doses as low as 0.2 mg/kg administered 24h before inoculation^40^. We therefore hypothesized that sufficient levels of sACE2_2_.v2.4-IgG1 would remain in the mice at the delivered dose. All mice pre-treated with a single-dose injection of sACE2_2_.v2.4-IgG1 before inoculation survived (**Figure 4D**) with slightly increased or no change in body weight (**Figure 4E**). The mice with single-dose of PBS injection before the inoculation all died at Day 7 and 8 with ∼30% loss of original body weight (**Figure 4D-E**). Furthermore, mice prophylactically administered sACE2_2_.v2.4-IgG1 showed no significant lung vascular leakage and edema at Day 7 post-inoculation, compared to marked increases in the control group (**Figure 4F-G**). Thus, single-dose sACE2_2_.v2.4-IgG1 pre-treatment prevented SARS-CoV-2-induced acute lung injury.

### Treatment with sACE2_2_.v2.4-IgG1 prevents lung vascular endothelial injury and ARDS and improves survival following inoculation with live SARS-CoV-2

To explore the therapeutic effects of sACE2_2_.v2.4-IgG1 on live SARS-CoV-2 infection, we chose two treatment timepoints: 12h or 24h following SARS-CoV-2 inoculation based on recent publications using neutralizing antibodies^40–43^. The timing of injection of sACE2_2_.v2.4-IgG1 post SARS-CoV-2 inoculation was chosen to begin treatment prior to onset of irreversible organ damage. Considering our mice were inoculated with high titer of SARS-CoV-2 (1×10^4^ PFU) and all died within 5-8 days of viral inoculation, initiation of treatment at 24 hours post viral inoculation may be equivalent to treating human patients at the onset of moderate to severe symptoms.

We used a 50% higher peptide dosage for the later time (24 hours) to increase the likelihood of efficacy in the face of prolonged viral replication. K18-hACE2 mice were all infected with SARS-CoV-2 isolate 1/2020 WA (1×10^4^ PFU) and then assigned to three treatment groups: PBS group (control), sACE2_2_.v2.4-IgG1 (10 mg/kg) treatment group (V2.4 12H) starting 12 hours post-inoculation, and sACE2_2_.v2.4-IgG1 (15 mg/kg) treatment group (V2.4 24H) starting 24 hours post-inoculation (**Figure 5A**). Treatment was administered IV daily for 7 days. Both, V2.4 12H and V2.4 24H treatment groups demonstrated markedly improved survival of 50%-60% at 2 weeks whereas all mice in the control treatment group died (**Figure 5B**). Initiation of treatment with sACE2_2_.v2.4-IgG1 at either 12h or 24h post viral inoculation also prevented weight loss (**Figure 5C**). We next determined efficacy of sACE2_2_.v2.4-IgG1 in preventing lung vascular endothelial injury and observed a marked reduction in lung transvascular albumin permeability (**Figure 5D-E**). However, edema formation was not changed by day 7 (**Figure 5F**). The viral load of lung tissues evaluated by real-time PCR (**Figure S4**) to detect the expression level of SARS-CoV-2 Spike and SARS-CoV-2 NSP, demonstrated near complete elimination of viral gene expression by Day 7 in both sACE2_2_.v2.4-IgG1 treatment groups (**Figure S4**).

**Figure 5.**
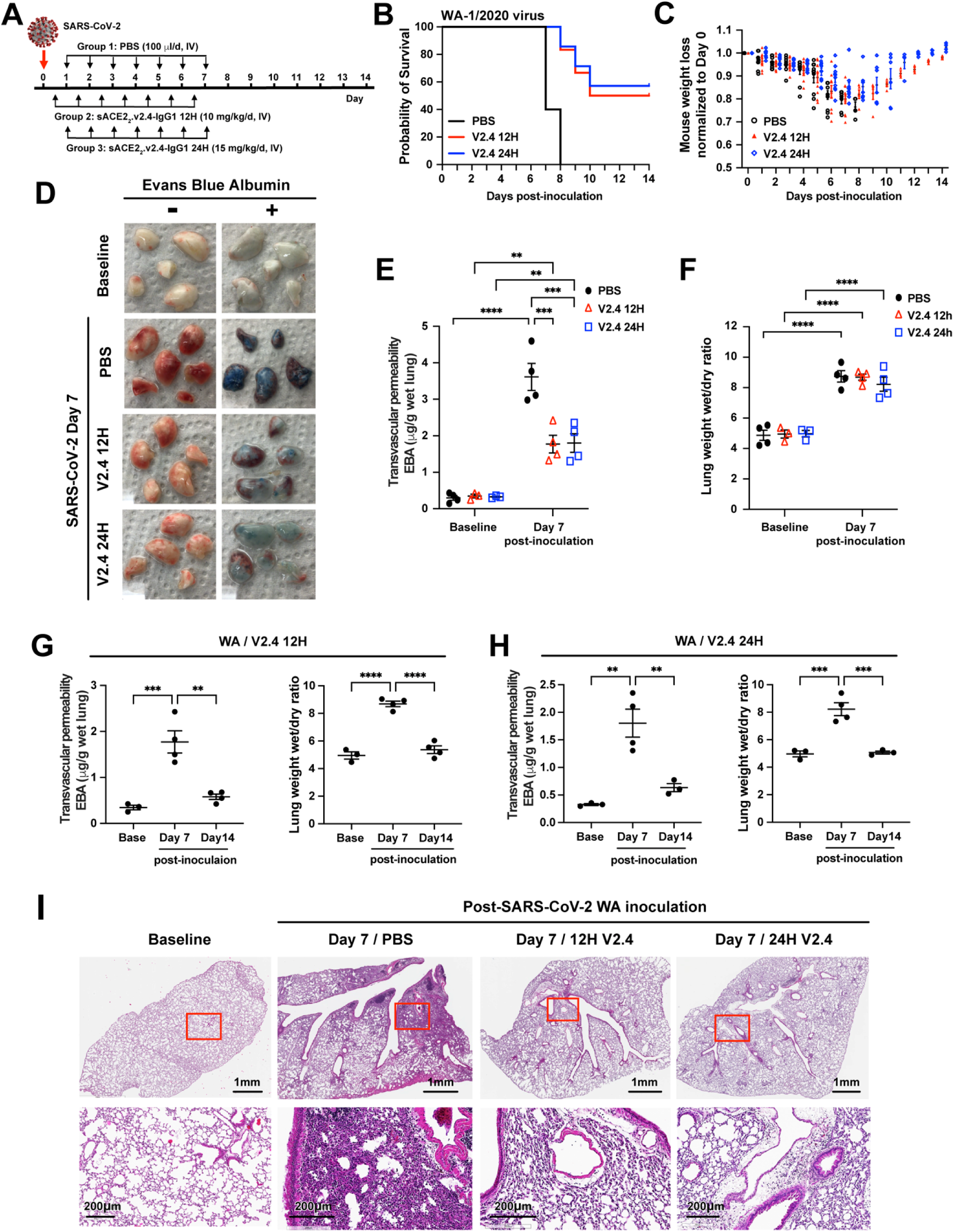
Treatment with sACE2_2_.v2.4 IgG1 mitigates lung vascular endothelial injury and ARDS and improves survival induced by live SARS-CoV-2 infection. (**A**) Experimental design to test the therapeutic efficacy of sACE2_2_.v2.4-IgG1. The K18 hACE2 transgenic mice were inoculated by SARS-CoV-2 isolate WA-1/2020 at 1×10^4^ PFU. Group 1 received control PBS via IV injection 24 hours post viral inoculation. Group 2 (V2.4 12H) received sACE2_2_.v2.4-IgG1 10mg/kg via IV injection 12 hours post inoculation, and then daily subsequent injections at the same dose. Group 3 (V2.4 24H) received sACE2_2_.v2.4-IgG1 15mg/kg via IV injection 24 hours post inoculation and then daily subsequent injections at the same dose. (**B**) Survival curves and (**C**) Weights for N = 10 mice for each group. (**D-F**) Mouse lungs were harvested at Day 7 post-inoculation for assessment of lung transvascular albumin permeability. (**D**) Macroscopic images of lungs at baseline and day 7 post-viral inoculation in the three experimental groups without EBA (Evans Blue Albumin) on the left and with EBA injection on the right. (**E**) Quantification of EBA in all three experimental groups. (**F**) Quantification of lung edema by wet/dry ratio in all three experimental groups were shown at baseline and Day 7 post-inoculation. (**G-H**) Time course of lung vascular permeability of Group 2 (V2.4 12H) (**G**) and Group 3 (V2.4 24H) (**H**) as assessed by the EBA assay and by the lung wet/dry ratio. (**I**) Representative H&E staining of lung sections at baseline (1^st^ column), control PBS group at day 7 post-inoculation with the WA isolate (2^nd^ column), sACE2.V2.4-IgG 12H treatment group at day 7 (3^rd^ column), and sACE2.V2.4-IgG 24H treatment group at day 7 (4^th^ column) post-inoculation with the WA isolate. The images in the first row are low magnifications. Rectangle areas (Red) are shown in higher magnification in the second row. Data are presented as mean ± SEM. **: P<0.01, ***: P<0.001, ****: P<0.0001 by Two-way ANOVA for **E** & **F**; one-way ANOVA for **G** & **H**.

As we did not observe any significant reduction in lung edema by Day 7 (**Figure 5F**) despite the marked improvement in survival (**Figure 5B**), we posited that reduction of edema may occur gradually in the surviving mice. We therefore assessed the time course of lung vascular permeability changes in both treatment groups at baseline, Day 7, and Day 14 post-inoculation with V2.4 12H and V2.4 24H treatments (**Figure 5G-H**). In the surviving mice (50-60% survival with peptide treatment versus 100% lethality in the no treatment group), lung transvascular albumin permeability and lung edema were restored to normal levels by Day 14 with either treatment (**Figure 5G-H**). Lung histology demonstrated that V2.4 12H and V2.4 24H treatment strikingly reduced immune cell filtration at Day 7 when compared to the control treatment group (**Figure 5I**). These results showed that treatment with engineered sACE2_2_.v2.4-IgG decoy peptide initiated as late as 24h after inoculation with a lethal dose of the SARS-CoV-2 WA-1/2020 isolate profoundly reduced lung vascular injury, prevented ARDS, and reduced mortality in a model of COVID-19.

### High affinity binding of sACE2_2_.v2.4 to S proteins of SARS-CoV-2 variants of concern

SARS-CoV-2 has a moderate mutation rate (10^-3^ substitutions per site per year ^44^) and variants have emerged with increased transmissibility and virulence, most notably the B.1.1.7 (Alpha), B.1.351 (Beta), P.1 (Gamma), and B.1.617.2 (Delta) VOCs. The variants have multiple mutations in S, including changes in immunodominant epitopes that cause partial immune escape ^11, 19, 20^ and a N501Y substitution within the RBD that increases monovalent affinity for ACE2 by 20-fold ^25^. Thus, we next assessed whether the decoy peptide was also efficacious against these new variants.

Using flow cytometry (a non-equilibrium experiment from which apparent binding strength was measured), binding of human cells expressing full-length trimeric S to wild type sACE2 and engineered sACE2.v2.4 was determined, using both monomeric sACE2 (residues 19-615) and dimeric sACE2_2_-IgG1 (ACE2 residues 19-732) to assess monovalent and avid binding, respectively (**Figure 6A**). Compared to wild type, monomeric sACE2.v2.4 showed markedly increased binding to all S proteins tested. The enhanced binding was less when avidity was considered, but nonetheless the engineered derivative remained tightly bound. Indeed, the engineered decoy receptor had increased binding signals to S from the newly circulating variants compared to the original SARS-CoV-2 isolate, consistent with some S variants showing tighter binding to ACE2. The level of sACE2 bound to S-expressing cells diminished at high concentrations, consistent with shedding of ACE2-bound S1. We explored this further by analysis of sACE2 binding to S variants from the B.1.429 (Epsilon), B.1.617.2 (Delta) and C.37 (Lambda) lineages (**Figure S5**). Again, monomeric sACE2.v2.4 bound considerably tighter than the wild type decoy, with smaller differences for sACE2_2_-IgG1 proteins where avidity masks affinity changes. Due to the decoy inducing S1 loss, detection at the cell surface of an N-terminal tag on S decreased.

**Figure 6.**
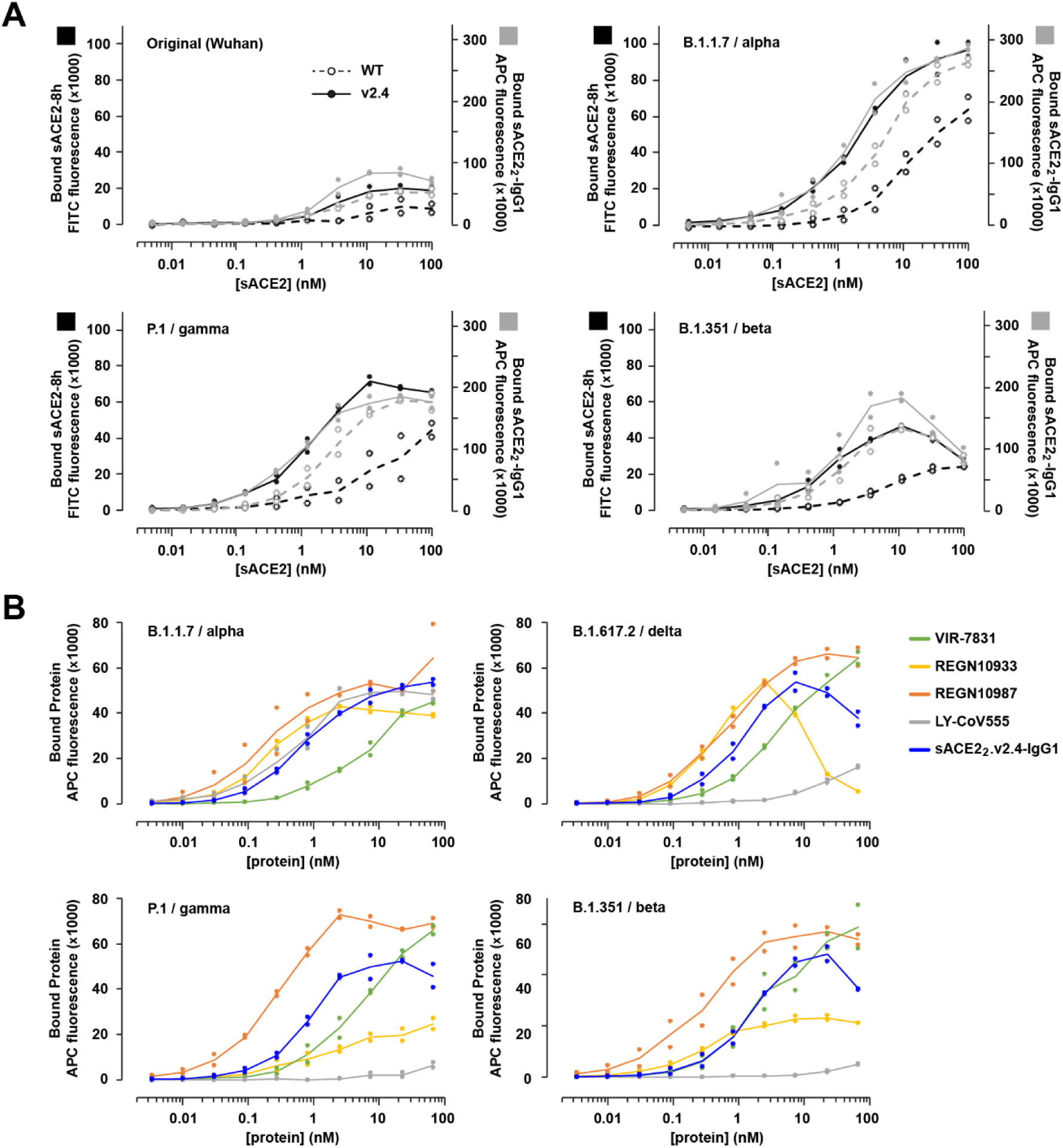
ACE2 decoys carrying v2.4 mutations bind S proteins from multiple highly transmissible SARS-CoV-2 VOCs. (**A**) Human Expi293F cells expressing myc-tagged S from four SARS-CoV-2 variants (Wuhan, B.1.1.7/alpha, P.1/gamma, and B1.351/beta) were incubated with monomeric sACE2-8h (black) or dimeric sACE2_2_-IgG1 (grey) and bound protein was detected by flow cytometry. N=2. (**B**) Binding of mAbs versus sACE2_2_.v2.4-IgG1 to S proteins of four VOCs (B.1.1.7/alpha, P.1/gamma, B1.351/beta, and B.1.617.2/delta) measured by flow cytometry. N=2.

Monoclonal antibodies in clinical development have shown mixed results for broad affinity against SARS-CoV-2 variants ^40^. Using flow cytometry, binding to S proteins from four VOCs was compared between sACE2_2_.v2.4-IgG1 and mAbs that have been used in the treatment of COVID-19 patients (**Figure 6B**): REGN10933 (casirivimab), REGN10987 (imdevimab), VIR-7831 (sotrovimab), and LY-CoV555 (bamlanivimab). In this assay, sACE2_2_.v2.4-IgG1 consistently shows high binding to S, better than VIR-7831 and LY-CoV555 but lower than REGN10987. The monoclonals REGN10933 and LY-CoV555 show markedly different binding strengths for the different S proteins. REGN10933 and sACE2_2_.v2.4-IgG1 engage the ACE2 interaction motif on S^45^ and both induce S1 shedding at high concentrations. Overall, sACE2_2_.v2.4-IgG1 broadly binds S proteins of SARS-CoV-2 VOCs and with comparable strength to mAbs that have shown efficacy in the clinic.

### sACE2_2_.v2.4-IgG1 prevents lung vascular endothelial injury, ARDS, and improves survival following inoculation with P.1 variant of concern

Based on our observations that sACE2_2_.v2.4-IgG1 binds S proteins of VOCs with even greater affinity than S protein of the original SARS-CoV-2 (**Figure 6** and **Figure S5**), we surmised that sACE2_2_.v2.4-IgG1 would also be therapeutically efficacious against VOCs. Here we tested the efficacy of sACE2_2_.v2.4-IgG1 against the P.1 variant, an aggressive and devastating SARS-CoV-2 variant in transmissibility and death in COVID-19 patients ^46^. We randomly assigned mice to three treatment groups: PBS group (control), sACE2_2_.v2.4-IgG1 (10 mg/kg) treatment group (V2.4 12H) starting 12 hours post-inoculation, and sACE2_2_.v2.4-IgG1 (15 mg/kg) treatment group (V2.4 24H) starting 24 hours post-inoculation. Early treatment with sACE2_2_.v2.4-IgG1 initiated at 12h post inoculation improved survival to 50-60% whereas delayed treatment initiation (starting at 24h post-inoculation) only delayed death but did not improve survival (**Figure 7A**). The mice in V2.4 12H treatment group also showed attenuated weight loss during the first 6 days following viral inoculation and the surviving mice gradually recovered their weight (**Figure 7B**). Both V2.4 12H and V2.4 24H treatment groups demonstrated marked reduction in lung transvascular albumin permeability (**Figure 7C**) and lung edema (**Figure 7D**) at Day 6, indicating mitigation of ARDS.

**Figure 7.**
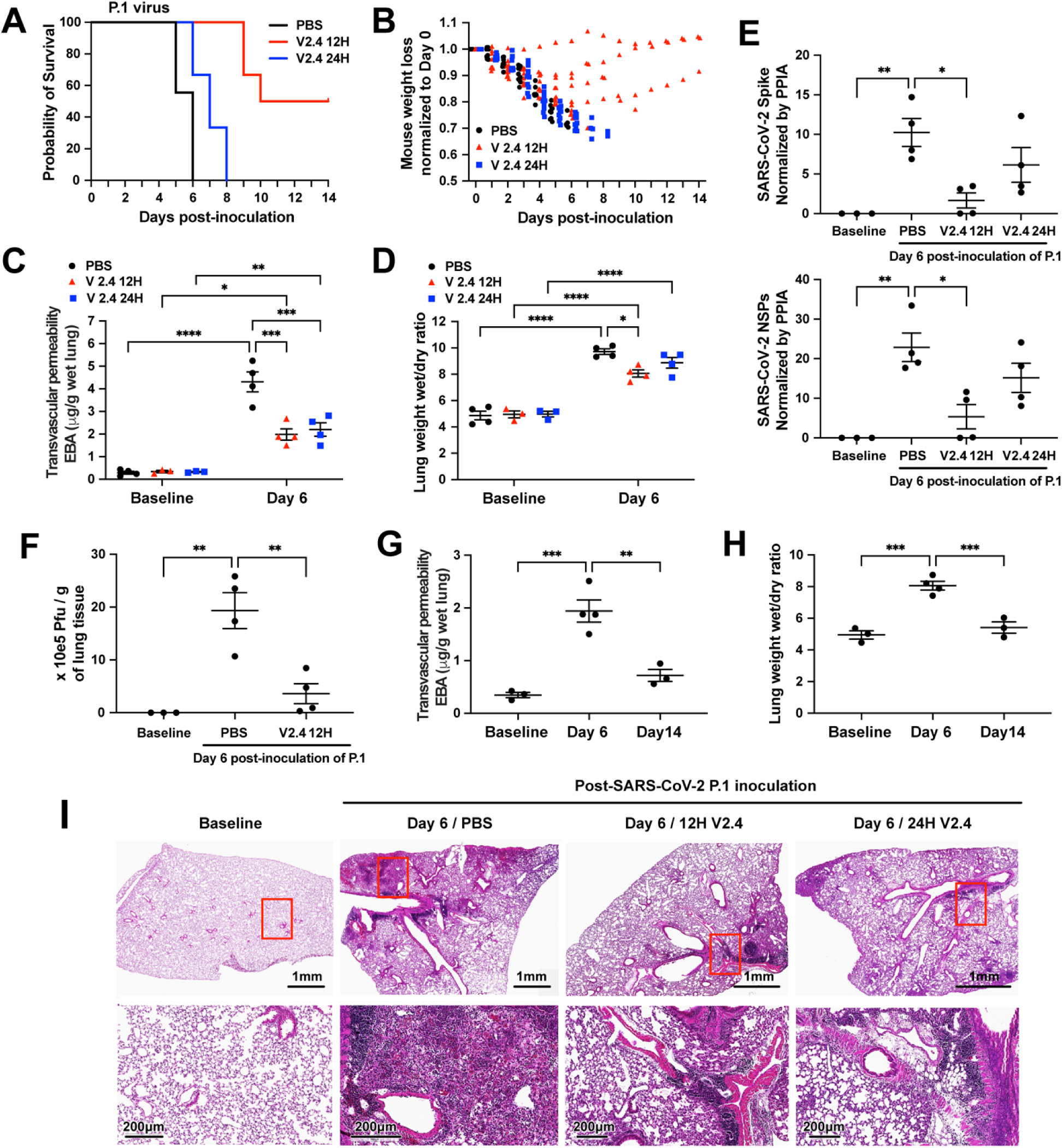
sACE2_2_.v2.4 IgG1 prevents lung vascular endothelial injury and ARDS and improves survival following infection with P.1 variant of concern. The three groups of K18 hACE2 transgenic mice were inoculated by SARS-CoV-2 variant P.1 (Brazil) at 1×10^4^ PFU. Group 1 (PBS): PBS was given by IV injection 24 hours post inoculation. Group 2 (V2.4 12H): sACE2_2_.v2.4-IgG1 10mg/kg were given by IV injection 12 hours post inoculation. Group 3 (V2.4 24H): sACE2_2_.v2.4-IgG1 15mg/kg were given by IV injection 24 hours post inoculation. The mice were injected once per day for 7 days. The survival probability was observed (**A**) and mouse weights were measured (**B**). N=10 for each group. (**C-D**) The mouse lungs were harvested at Day 6 post-inoculation to evaluate lung transvascular permeability – EBA assay (**C**) and lung wet/dry ratio (**D**) with baseline mouse lungs as control. (**E**) The viral loads of SARS-CoV-2 in the lungs harvested at baseline and Day 6 post-inoculation of SARS-CoV-2 variant P.1 (Brazil) were measured by real-time quantitative PCR for the mRNA expression level of SARS-CoV-2 Spike and SARS-CoV-2 NSPs. (**F**) Viral Plague Form Assay was performed to measure the viral loads of SARS-CoV-2 in the lungs harvested at baseline and Day 6 post-inoculation of SARS-CoV-2 variant P.1 (Brazil). (**G-H**) Time course of lung transvascular permeability of V2.4 12H treatment group. The EBA assay (**G**) and Wet/dry ratio (**H**) were measured at baseline, Day 6 and Day 14 post-inoculation. (**I**) Representative H&E staining of lung sections at baseline (1^st^ column), control PBS group at day 7 post-inoculation with the P.1 variant (2^nd^ column), sACE2.V2.4-IgG 12H treatment group at day 7 (3^rd^ column), and sACE2.V2.4-IgG 24H treatment group at day 7 (4^th^ column) post-inoculation with the P.1 variant. The images in the first row are low magnifications. Highlighted areas (Red) are shown in higher magnification in the second row. N=4 for **C, D, E, F, G**, and **H**. Data are presented by mean ± SEM. *: P<0.05, **: P<0.01, ***: P<0.001, ****: P<0.0001 by Two-way ANOVA for **C** & **D**; one-way ANOVA for **E, F, G** & **H**.

The viral load of lung tissues evaluated by real-time PCR showed that initiating early treatment with sACE2_2_.v2.4-IgG1 reduced the viral copy number by 75%, whereas delayed treatment initiation at 24h only reduced viral gene copies by 50% (**Figure 7E**). Using the viral plaque forming assay in lung tissue as a functional measure of viral load in lungs, we found that early sACE2_2_.v2.4-IgG1 treatment markedly reduced viral plaque forming capacity on Day 6 by over 80% (**Figure 7F**).

To study robustness of recovery of lung vascular function in mice that survived inoculation with the P.1 variant following sACE2_2_.v2.4-IgG1 treatment, we assessed lung transvascular albumin permeability at baseline, Day 6, and Day 14 post-inoculation and with early V2.4 12H treatment (**Figure 7G-H**). We observed that lung transvascular albumin permeability returned to near-baseline levels in surviving mice by Day 14 (**Figure 7G-H**). Histological analysis demonstrated reduced immune cell infiltration following infection with P.1 variant at Day 7 in the V2.4 12 hours and V2.4 24 hours treatment groups (**Figure 7I**). Thus, sACE2_2_.v2.4-IgG1 decoy peptide proved to be more effective when administered early during the infection with P.1 variant of concern.

## DISCUSSION

Acute lung injury and its progression to the severe acute respiratory distress syndrome (ARDS) results in respiratory failure due to protein-rich lung edema formation. ARDS is the leading cause of death in COVID-19 patients ^47^. The severe lung edema is the consequence of breakdown of the lung endothelial barrier ^29, 30^ due to injury of endothelial adherens junction, loss of lung endothelial cells as a consequence of cell death and denudation ^28, 48^, and suppression of endothelial cell regeneration pathways ^49^. While there is growing evidence of endothelial cell disruption in autopsy samples of COVID-19 patients ^50, 51^, studies have focused on factors such as pro-thrombotic transition of the lung endothelium ^52^. Our studies showed loss of VE-cadherin and lung endothelial cell injury in the humanized K18-hACE2 model of COVID-19. Endothelial injury as quantified by measuring lung vascular endothelial permeability and resulting pulmonary edema and ARDS were severe in mice inoculated with SARS-CoV-2 isolate 1/2020 WA, and were even more pronounced after inoculating the mice with the P.1 VOC. The more severe lung vascular injury explains the devastatingly high mortality rates in patients infected with the new more toxic variants ^44, 46^. We also demonstrated higher affinity binding of ACE2 to Spike proteins of SARS-CoV-2 VOCs as compared to SARS-CoV-2 isolate 1/2020 WA.

The transgenic K18-hACE2 mice used to study the responses to SARS-CoV-2 virus (both of the original SARS-CoV-2 isolate 1/2020 WA and the P.1 VOC) similarly expressed hACE2 in lung alveolar epithelial type II cells ^27^. Thus, lung vascular injury induced by SARS-CoV-2 can only be ascribed to virus binding and entry into type II cells. The augmented endothelial injury and higher mortality with the P.1 variant was consistent with greater binding efficiency of ACE2 to the Spike protein of the variant. Lung endothelial injury is due to release of inflammatory cytokines or damage associated molecular pattern (DAMP) molecules brought about by the infected type II alveolar epithelial cells ^53^. A key factor in the pathogenesis of lung endothelial injury is that the lung cells uniquely express higher levels of pro-inflammatory genes compared to endothelia of other organs ^54^. This is believed to reflect the key role of the lung endothelium in the host-defense function of lungs (61). The enrichment of inflammatory genes in lung endothelial cells may therefore increase the vulnerability of the lung endothelial barrier cells to the “cytokine storm” elicited by COVID-19 infection^53^. Thus, augmentation of the cytokine storm by the P.1 variant may have resulted in more severe lung endothelial injury and edema and mortality as compared to SARS-CoV-2 isolate 1/2020 WA.

The development of mRNA-based and adenoviral vaccines to induce immunity against SARS-CoV-2 with reported efficacy rates of 70% to 95% ^1–7^ has been a great success in translational science. However, the growing number of mutations in SARS-CoV-2 Spike protein has resulted in the emergence of multiple SARS-CoV-2 VOCs characterized by high transmissibility and evasion to varying degrees of the humoral immune response ^9, 11, 55–63^. The high transmissibility of these variants as a result of antigenic drift is due in part to the higher binding affinity of the Spike protein for ACE2, resulting in greater cell entry and infectivity ^59, 64^. However, we showed that the higher affinity of variant Spike proteins for ACE2 can be therapeutically leveraged to engineer decoy peptides based on ACE2 that compete for Spike protein binding. We developed and tested the effectiveness of sACE2_2_.v2.4-IgG1 peptide as an optimized receptor decoy. The decoy peptide acted as a ‘sponge’ for SARS-CoV-2 and prevented infection by blocking binding to ACE2-expressing cells in the lung.

While the emergence of SARS-CoV-2 variants of concern reduced the efficacy of vaccines and therapeutic neutralizing antibodies ^11, 58^, our data demonstrate that the engineered sACE2_2_.v2.4-IgG1 peptide has even greater affinity for Spike variants. Previous studies have engineered high-affinity sACE2 decoys, some achieving even tighter affinity than sACE2_2_.v2.4 by combining more mutations ^22, 65^, yet minimizing the number of substitutions ensures that the decoy receptor closely resembles native ACE2, reducing opportunities for viral escape or potential drug immunogenicity.

We showed prophylactic administration of sACE2_2_.v2.4-IgG1 fully protected against SARS-CoV-2-induced acute lung injury. More importantly, we demonstrated that sACE2_2_.v2.4-IgG1 treatment as late as 24h or 12h after virus inoculation reduced mortality by 50-60% for the WA-1/2020 and P.1 viruses, respectively. While treatment at 24h was too late to prevent death from the more virulent P.1 virus, lung vascular leakage, pathology and viral load were nonetheless reduced with survival time extended, demonstrating efficacy at reducing symptoms. While it is difficult to directly model the time course of human COVID-19 in animal studies, our therapeutic regimen compares exceedingly well to parallel studies using neutralizing monoclonal antibodies (see **Table S1 and S2** for comparisons of prophylaxis and therapy studies). Indeed, our use of a highly lethal infection model permitted broader assessment of how the peptide mitigates lung injury and death than many other studies in animal models that show minimal disease. A comparable assessment of monoclonal antibodies that are in the clinic demonstrated efficacy when the proteins were administered at 24h post inoculation in K18-hACE2 mice, but animals were infected at lower SARS-CoV-2 titers (1×10^3^ FFU) and weight loss mostly began at day 3 or 4^40^. In our model, mice were inoculated with high virus titer (1×10^4^ PFU), began to lose weight early by day 2, and all died within 5-7 days. We reason that initiation of treatment at 24h post viral exposure in this model may be equivalent to treating human patients at the onset of moderate to severe symptoms.

Although we focused on using intravenous delivery for our efficacy studies, the pharmacokinetic analysis demonstrated that inhaled or intratracheally applied peptide resulted in stable lung tissue concentrations. Therefore, for translational studies, one would envision staged therapy in which sACE2_2_.v2.4-IgG1 is delivered locally in lungs via inhalation during the early post-exposure period and then systemically via intravenous route during later stages after virus dissemination.

We found that sACE2_2_.v2.4-IgG1 was effective in sharply reducing lung vascular injury, ARDS, and mortality in mice infected with the P.1 variant, among the most highly transmissible VOCs with high mortality rate ^19, 44, 46, 64^. We also showed that the peptide bound all variant Spike proteins with similar or even greater affinity than the original SARS-CoV-2 isolate. This was also the case with the recent B.1.617.2 (Delta) virus, now the primary cause of morbidity and mortality world-wide. Based on these data, it is reasonable to conclude that the *in vivo* efficacy will be similar for distinct variants.

We observed that the P.1. variant required earlier initiation of treatment (at 12h instead of 24h) following virus exposure to achieve 50-60% reduction in mortality. However, because the binding affinity for the P.1. Spike protein was even greater than the original SARS-CoV-2 isolate, the need for earlier treatment initiation likely reflected rapid dissemination and replication of the P.1. variant prior to therapy initiation. In this context, it would be useful to address whether efficacy improves by using either a higher peptide dose or increased administration frequency when initiating the delayed treatment against variants with high infectivity and replication rates.

Broad affinity of sACE2_2_.v2.4-IgG1 against SARS-CoV-2 variants as well as other coronaviruses that use ACE2 as an entry receptor^25^ is an advantageous feature compared to most mAbs where cocktails are required^66^. We also showed that sACE2_2_.v2.4-IgG1 reduced S1 at a cell membrane, suggesting that it might irreversibly inactivate the virus through S1 shedding. ACE2-catalyzed turnover of angiotensin and kinin peptide hormones can also directly prevent ARDS ^23, 67^. These multiple mechanisms of action provide sACE2_2_.v2.4-IgG1 with unique characteristics compared to mAbs in the clinic.

While we demonstrated the *in vivo* efficacy of the engineered sACE2_2_.v2.4-IgG1 peptide against two distinct live SARS-CoV-2 variants in the K18-hACE2 model, a non-human primate model ^52^ may mirror aspects of human COVID-19 more accurately, as it would allow for SARS-CoV-2 infection not only of the epithelium but also of other cell types such as the endothelium, which also express ACE2 and TMPRSS2, and can be infected by SARS-CoV-2 (**Figure 4**). Another limitation is our focus on intravenous delivery, whereas intratracheal or inhalation delivery may be more readily applied in patients, especially in the outpatient setting. It would be useful in future studies to use varying doses and delivery route combinations for defined disease stages to identify the optimal combination prior to initiating human efficacy trials. Finally, we chose to fuse the ACE2 peptide to an unmodified Fc from IgG1 (isoallotype nG1m1), whereas others have considered fusions to Fc mutants or IgG4 to dampen interactions with FcγR subtypes that might contribute to inflammation^24, 68^. We instead reasoned that IgG1 effector functions will be necessary for optimum *in vivo* protection.

In summary, we show for the first time that an engineered decoy ACE2 peptide demonstrating much higher affinity for the SARS-CoV-2 Spike protein is efficacious *in vivo* against multiple SARS-CoV-2 variants. This peptide prevented viral entry into cells and the peptide’s efficacy against a variant of concern prevented lung endothelial injury and ARDS and significantly reduced mortality. The results show the potential of this engineered peptide to treat COVID-19 patients and others with inadequate antibody titer to protect against emerging, more virulent SARS-CoV-2 variants.

## METHODS

### SARS-CoV-2 viruses and cells used for viral replication

2019n-CoV/USA_WA1/2020 isolate of SARS-CoV-2 (NR-52281) and SARS-CoV-2 Isolate hCoV-19/Japan/TY7-503/2021 (Brazil P.1) (NR-54982) were obtained from BEI Resources, NIAID, NIH. Vero E6 (CRL-1586, American Type Culture Collection, ATCC) were cultured at 37°C in Dulbecco’s Modified Eagle medium (DMEM) supplemented with 10% fetal bovine serum (FBS), 1 mM sodium pyruvate, 1× non-essential amino acids, and 100 U/ml of penicillin–streptomycin. The SARS-CoV-2 were propagated in Vero E6 cells (CRL-1586; American Type Culture Collection). The supernatant was collected upon observation of cytopathic effect. The debris was removed by centrifugation and passage through a 0.22 μm filter. Supernatant was then aliquoted and stored at −80°C. The virus titers were quantitated by a plaque forming assay using Vero E6 cells.

### Inoculation of SARS-CoV-2 in K18-hACE2 transgenic mice and administration of sACE2_2_.v2.4

Biosafety level 3 (BSL-3) protocols for animal experiments with live SARS-CoV-2 were performed by personnel equipped with powered air-purifying respirator in strict compliance with the National Institutes of Health guidelines for humane treatment and approved by the University of Illinois Animal Care & Use Committee (UIC protocol 18-076). Hemizygous K18-hACE mice with c57BL/6J background (strain#034860: B6.Cg-Tg(K18-ACE2)2Prlmn/J) were purchased from The Jackson Laboratory. Animals were housed in groups and fed standard chow diets. The 8-10 weeks old mice were anesthetized by ketamine/xylazine (50/5 mg/kg, intraperitonially). The mice were then inoculated intranasally with 1×10^5^ PFU (plaque-forming units) of SARS-CoV-2 isolate WA or 1×10^4^ PFU of SARS-CoV-2 isolate (WA) or the P.1 variant suspended in 20 μL of sterile phosphate-buffered saline. sACE2_2_.v2.4 was administered intravenously (retro-orbital injection) either 12 hours before virus inoculation (single-dose) or 12 hours / 24 hours post virus inoculation (once a day for continuous 7 days) to the anesthetized K18-hACE2 transgenic mice. All efforts were made to minimize animal suffering. Genotyping of mice was performed by PCR using tail DNA. For the number of animals needed to achieve statistically significant results, we conducted a priori power analysis. We calculated power and sample sizes according to data from small pilot experiments, variations within each group of data, and variance similarities between the groups that were statistically compared. Animals with sex- and age-matched littermates were randomly included in experiments. No animals had to be excluded attributed to illness after experiments. Animal experiments were carried out in a blinded fashion whenever feasible.

### Plaque forming assay

Vero E6 (CRL-1586, ATCC) were seeded at a density of 2.5×10^5^ cells per well in flat-bottom 12-well tissue culture plates. The following day, media was removed and replaced with 200 μL of 10-fold serial dilutions of the material to be tittered, diluted in DMEM+2% FBS. One hours later, 1 mL of 0.8% agarose overlay was added. Plates were incubated for 72 hours, then fixed with 4% paraformaldehyde (final concentration) in phosphate-buffered saline for 30 minutes. Plates were stained with 0.05% (w/v) crystal violet in 20% methanol and washed twice with distilled, deionized H_2_O.

### Generation of pseudoviruses

Recombinant Indiana vesicular stomatitis virus (rVSV) expressing the SARS-CoV-2 spike was generated as previously described^69^. HEK293T cells were grown to 80% confluency before transfection with pCMV3-SARS-CoV-2-spike (Sino Biological) using Lipofectamine®3000 (Invitrogen). Cells were cultured overnight at 37□°C with 5% CO_2_. The next day, medium was removed and VSV-G pseudotyped ΔG-luciferase (a gift from Dr. David D. Ho, Columbia University) was used to infect the cells in DMEM at a MOI of 3 for 2 h. After infection, cells were washed with PBS twice, and new complete DMEM culture medium was added. DMEM supplemented with 2% fetal bovine serum, 100 IU/ml of penicillin and 100 μg/ml of streptomycin. The supernatant were harvested and clarified by centrifugation at 800*g* for 10 min, and filtered by 0.45-μm pore size (Millipore) before aliquoting and storing at −80□°C. Prior to infection of target cells, incubate the viral stock with 20% I1 hybridoma (anti-VSV-G) supernatant (ATCC; CRL-2700) for 1 h at 37°C to neutralize contaminating G*ΔG rVSV-luciferase particles.

### Virus entry evaluation by luciferase assay

For *in vitro* studies, cultured cells were preincubated with 5 and 25 µg/ml soluble sACE2_2_-IgG1 (wt) and sACE2_2_.v2.4-IgG1 variant for 1 hour. The SARS-CoV-2 pseudovirus (MOI=0.1) was then added to the cultured cells. The cells were harvested for luciferase assay after 24 hours. For *in vivo* studies, 10 mg/kg sACE2_2_.v2.4-IgG1 (wt), sACE2_2_.v2.4-IgG1 and control peptide buffer were intravenously administrated into K-18 hACE-2 mice for 30 minutes prior to SARS-CoV-2 pseudo-entry virus (10^6^ pfu) i.p. injection for 1 dpi. Tissue lysates were prepared, and virus entry ability in the selected organs was evaluated by luciferase activity using the luciferase assay system (Promega) and a microplate luminometer. Data are mean ± SEM of n = 4 replicates.

### RNA isolation and quantitative real-time PCR

RNA is extracted from homogenized lung tissues using TRIzol™ Reagent (ThermoFisher #15596026) according to the manufacturer’s protocol. Then RNA is quantified by Nanodrop 1000 (ThermoFisher) and reverse transcribed into cDNA using High-Capacity cDNA Reverse Transcription Kit (ThermoFisher #4368814). FastStart Universal SYBR Green Master (ThermoFisher # 4913850001) was used for relative quantification of cDNA on the ViiA 7 Real-Time PCR System (ThermoFisher) (Primer information included in **Table S3**).

### Measurement of viral burden

Tissues were weighed and homogenized in 1 ml Trizol solution. Tissue homogenates were clarified by centrifugation at 10,000 rpm for 5 min and stored at −80°C. RNA was extracted following a Trizol protocol. RNA was reverse-transcribed with Superscript III (Invitrogen) using random primers. Synthesized cDNA samples were amplified in the ABI PRISM 7000 Sequence Detection System (Applied Biosystems) thermocycler using SYBR Green using primers designed to target a highly conserved region of the Spike gene and NSP gene (Primer information included in **Table S4**).

### Reagents

Polyclonal anti-SARS-CoV-2 spike glycoprotein was obtained from BEI Resources. Luciferase assay system (E1500) was purchased from Promega. Enhanced chemiluminescence (ECL) Western blotting Detection Reagents and nitrocellulose membranes (Hybond-ECL) were from Amersham Biosciences Corp. Lipofectamine® 3000 transfection reagents were from Invitrogen. Antibodies for flow cytometry are described elsewhere in Methods.

### Lung vascular permeability measurements in mice

The Evans Blue–Albumin pulmonary transvascular flux measurements were performed to measure lung vascular leakage. Briefly, 200 μL Evans blue–albumin (1% Evans blue dye [EBD] and 4% albumin in PBS) was administered intravenously (retro-orbital injection) into anesthetized mice and allowed to circulate in the blood vessels for 30 minutes. Mice were scarified and lungs were perfused by 10 mL PBS. Lung tissues were then excised, weighed, homogenized in 1 mL PBS and extracted overnight in 2 mL formamide at 60°C. Samples were centrifuged at 10000g for 5 minutes. Evans blue concentration in lung homogenate supernatants was quantified by the spectrophotometric method at absorbance of 620 nm. Tissue EBD content (μg EBD/g fresh lung tissue) was calculated by comparing tissue supernatant A_620_ readings with an EBD standard curve. Concentration of Evans blue dye was determined in micrograms per gram of wet lung tissue. The ratio of wet lung to dry lung weight for edema measurement was calculated.

### Histology and imaging

Animals were euthanized before harvest and fixation of tissues. Lung lobes were fixed with 4% PFA (Paraformaldehyde) for 48 hours before further processing. Tissues were embedded in paraffin, and sections were stained with hematoxylin and eosin, Trichrome blue and Sirius Red for collagen deposition. Images were taken with a Zeiss microscope and analyzed by Zen software (Zeiss).

### Mouse cell isolation

The mouse cell isolation was described previously ^28^. Briefly, flushed mouse lung tissue was minced and digested with 3 ml Type 1 collagenase I (2 mg/ml in PBS) at 37°C water bath for 1 hour. Mixtures were titrated with #18 gauge needles and then pipetted through a 40 μm disposable cell strainer. After centrifuging 300 g for 5 minutes and washing with PBS, isolated cells were treated with red blood cell lysis buffer (eBioscience) for 5 minutes on ice to lyse red blood cells.

### Flow cytometry

After isolation, remaining cells were incubated with anti-mouse CD16/CD32 (1:50, BD Pharmingen#553142) to block endogenous Fc for 10 minutes on ice. After this, cells were stained with antibodies including CD45-EF450 (1:2000, eBioscience #48-0451-82) and CD31-APC (1:100, eBioscience #17-0311-82) for 45 minutes at 4°C. After wash, the cells were resuspended in 500 μl buffer and analyzed on an LSR Fortessa (BD Pharmingen) cell analyzer. Obtained data were analyzed by Summit software (Beckman Coulter).

### Cell culture

The 2019n-CoV/USA_WA1/2019 isolate as well as the P.1 variant of SARS-CoV-2 and ACE-2 expressing A549 cell lines were obtained from BEI Resources. Vero E6 (CRL-1586; American Type Culture Collection) Vero CCL81 (American Type Culture Collection), ACE-2 expressing A549 cells and standard A549 cells were cultured at 37□ °C in Dulbecco’s modified Eagle’s medium (DMEM) supplemented with 10% fetal bovine serum (FBS), 10□mM HEPES (pH□7.3), 1□mM sodium pyruvate, 1× non-essential amino acids and penicillin–streptomycin. Primary human lung microvascular endothelial cells (hLMVECs) from Lonza were cultured in EBM-2 supplemented with 10% endotoxin-free fetal bovine serum (Omega Scientific). Infectious stocks were grown by inoculating Vero CCL81 cells and collecting supernatant upon observation of cytopathic effect; debris were removed by centrifugation and passage through a 0.22-μm filter. Supernatant was then aliquoted and stored at −80□°C.

Expi293F cells (Thermo Fisher Scientific) were cultured in Expi293 Expression Medium (Thermo Fisher Scientific) at 37 °C, 125 rpm, 8% CO_2_. Expi293F cells were transfected at a density of 2 × 10^6^ /ml with 400 ng (pCEP4-myc-ACE2 plasmids) or 500 ng (pCEP4-myc-S plasmids) per ml of culture using ExpiFectamine 293 (Thermo Fisher Scientific) according to the manufacturer’s directions. For in vitro binding assays using flow cytometry, transfection enhancers were excluded and cells were harvested 24-30 h post-transfection. For production of monomeric sACE2-8h, ExpiFectamine 293 transfection enhancers (Thermo Fisher Scientific) were added ∼20 h post-transfection and media was collected for protein purification 4 to 6 days later.

Dimeric sACE2_2_-IgG1 proteins were expressed in ExpiCHO-S cells (Thermo Fisher Scientific) cultured in ExpiCHO Expression Medium (Thermo Fisher Scientific) at 37 °C, 125 rpm, 8% CO_2_. The cells were transfected with ExpiFectamine CHO (Thermo Fisher Scientific) according to the manufacturer’s directions, using 500-1000 ng plasmid per ml of culture. ExpiFectamine CHO Enhancer (6 μl per ml of culture) and ExpiCHO Feed (240 μl per ml of culture) (Thermo Fisher Scientific) were added 18-22 hours post-transfection and the temperature was decreased to 33 °C. At day 5 post-transfection, ExpiCHO Feed (240 μl per ml of culture) was again added. On days 9 and 11, the CO_2_ was stepped down to 7% and 6%, respectively. The media was harvested on days 12 to 14 for protein purification.

### Plasmids

Residue numbers for proteins begin with amino acid (a.a.) 1 as the start methionine. Plasmids for the expression of myc-tagged human ACE2 (pCEP4-myc-ACE2, Addgene No. 141185), 8his-tagged monomeric sACE2 (ACE2 a.a. 1-615; pcDNA3-sACE2(WT)-8his, Addgene No. 149268, and pcDNA3-sACE2v2.4-8his, Addgene No. 149664), and human IgG1-Fc fused dimeric sACE2_2_ (ACE2 a.a. 1-732; pcDNA3-sACE2-WT(732)-IgG1, Addgene No. 154104, and pcDNA3-sACE2v2.4(732)-IgG1, Addgene No. 154106) are previously described. Human codon-optimized S (GenBank No. YP_009724390.1) was subcloned into pCEP4 (Invitrogen) from pUC57-2019-nCoV-S(Human) (distributed by Molecular Cloud on behalf of Haisheng Yu, Chinese Academy of Medical Sciences) with a N-terminal HA leader (MKTIIALSYIFCLVFA), myc-tag (EQKLISEEDL), and linker (GSPGGA) upstream of the mature polypeptide (a.a. V16-T1273). All mutations were made by extension overlap PCR and confirmed by Sanger sequencing (ACGT, Inc.) Compared to the SARS-CoV-2 Spike reference sequence (GenBank Accession YP_009724390.1), the variant myc-tagged S proteins cloned here have the following mutations: P.1 (L18F, T20N, P26S, D138Y, R190S, K417T, E484K, N501Y, D614G, H655Y, T1027I), B.1.351 (L18F, D80A, D215G, ΔL242-L244, K417N, E484K, N501Y, D614G, A701V), B.1.1.7 (ΔH69-V70, ΔY144, N501Y, A570D, D614G, P681H, T716I, S982A, D1118H), B.1.429 (W152C, L452R, D614G), and B.1.617.2 (T19R, G142D, A222V, L452R, T478K, D614G, P681R).

### Assessment of ACE2 mutants

Expi293F cells transfected with pCEP4-myc-ACE2 plasmids were collected 24 h post-transfection (600 × g, 60 s), washed with ice-cold Dulbecco’s phosphate buffered saline (PBS) containing 0.2% bovine serum albumin (BSA), and stained with 1:50 RBD-sfGFP expression medium (prepared as previously described ^22^ and 1:250 anti-myc Alexa 647 (clone 9B11, Cell Signaling Technology) in PBS-BSA. After 30 minutes incubation on ice, cells were washed twice in PBS-BSA and analyzed on a BD Accuri C6 using instrument software. Gates were drawn around the main population based on forward and side scatter properties, followed by a gate for medium Alexa 647 fluorescence to control for any expression differences between mutants (**Figure S6A-B** and as previously described ^22^). Mean sfGFP fluorescence was measured and background fluorescence of negative cells was subtracted. All replicates were from independent, distinct samples.

### Purification of sACE2_2_-IgG1

ExpiCHO-S expression medium was centrifuged (800 × g, 4 °C, 10 minutes) to remove cells. 1M Tris base was added to the supernatant until pH ∼7.5. The medium was centrifuged again (15,000 × g, 4 °C, 20 minutes). KanCapA resin (Kaneka Corporation, equilibrated in PBS) was incubated with the supernatant (2 ml resin per 100 ml medium) for 2 h while the sample was rotated at 4 °C. The resin was collected by passing the sample through an empty chromatography column and was washed with 10 column volumes (CV) Dulbecco’s phosphate buffered saline (PBS). Protein was eluted with 4 CV 60 mM sodium acetate pH 3.7 into a vessel containing 2 CV 1M Tris pH 8.0, yielding a final pH ∼6. The pH was raised to 7 by the addition of 1M Tris base (1 to 2 CV). The eluate was concentrated with a 50 kD MW cutoff (MWCO) centrifugal filtration unit (Millipore) and injected on a HiLoad Superdex 200 pg 16/600 gel filtration column (GE Healthcare) with PBS as the running buffer. Peak fractions at the expected MW of a dimer were pooled, concentrated (> 30 mg/ml), and aliquots were frozen in liquid nitrogen and stored at -80 °C. Purity was routinely > 98% based on SDS polyacrylamide gel electrophoresis (SDS-PAGE). Concentrations were determined by absorbance at 280 nm using the calculated molar extinction coefficient for the mature, monomeric polypeptide. For mouse studies, protein aliquots were thawed, diluted to 5 mg/ml with PBS, and filter (0.2 μm) sterilized ready for use.

### Purification of sACE2-8h

Expi293F expression medium was centrifuged to remove cells (800 × g, 4 °C, 10 minutes) and then to remove smaller particulates (15,000 × g, 4 °C, 20 minutes). HisPur Ni-NTA resin (Thermo Fisher Scientific) equilibrated in PBS was added (0.5 ml resin per 100 ml medium). The sample was rotated at 4 °C, 90 minutes. The resin was collected by passing through an empty chromatography column and then washed with >20 CV PBS. Proteins were eluted using a step gradient of PBS containing 20 mM, 50 mM, and 250 mM imidazole (pH 8.0, 12 CV for each fraction). The 50 and 250 mM imidazole fractions contained pure protein based on Coomassie-stained SDS-PAGE and were pooled and concentrated using a 30 kD MWCO centrifugal filtration unit (Millipore). Proteins were separated on a Superdex 200 Increase 10/300 GL column (GE Healthcare) with PBS as the running buffer. Peak fractions at the expected MW of a monomer were pooled, concentrated (∼5 mg/ml), and aliquots were snap frozen and stored at -80 °C. Concentrations were based on UV (280 nm) absorbance using the molar extinction coefficient for the mature, monomeric polypeptide.

### Toxicology

Animal experimental procedures were reviewed and approved by the University of Illinois Institutional Animal Care and Use Committee (UIUC protocol 20127). Toxicity of each peptide was assessed in 5 female and 5 male CD-1 IGS mice (8 weeks old). For sACE2_2_.v2.4 without any tags or fusion partner (corresponding to ACE2 residues 19-732, protein provided by Orthogonal Biologics, Inc.), mice were IV administered (tail vein, 0.5 mg/kg) protein twice daily for 5 consecutive days (days 0-4) and sacrificed day 7. Multiple doses were administered due to the anticipated short serum half-life. To assess toxicity of sACE2_2_.v2.4-IgG1, a single protein dose was IV administered (tail vein, 2.0 mg/kg) and mice were sacrificed on day 7. Toxicity was assessed by body weight, complete blood count, serum chemistry, and necropsy/histopathology. No differences were observed compared to vehicle treated mice.

### Pharmacokinetics

8-week-old CD-1 IGS mice (3 female and 3 male per time point) were IV administered protein solutions. Mice were sacrificed and blood was drawn for serum analysis at designated time points. sACE2_2_.v2.4 (without any tags or fusion partner) and sACE2_2_.v2.4-IgG1 were injected via the tail vein at doses of 0.5 mg/kg and 2.0 mg/kg, respectively. Catalytic activity in serum was measured using the Fluorometric ACE2 Activity Assay Kit (BioVision), sACE2 levels were measured by Human ACE-2 ELISA Kit (RayBiotech), and levels of the IgG1 Fc moiety were measured by Human IgG ELISA Kit (Immunology Consultants Laboratory). To characterize different routes side-by-side, peptides were administered to C57BL/6 mice, 3 males per time point. Animals were 25-35 g and 8-10 weeks old. Administration was via: tail vein injection (2.0 mg peptide / kg); intubation-mediated intratracheal (IT) instillation (1.0 mg peptide / kg); or inhalation where peptide solution was nebulized into the animal holding chamber. Peptide solutions were provided sterile-filtered in PBS at 0.4 mg/ml (IV), 1.0 mg/ml (IT), or 50 mg/ml (inhalation). For IT dek, mice were anaesthetized with 2-3% isoflurane/oxygen and peptide solution (50 µL) / air (150 µL) was delivered with a blunt 20 G catheter. At time points, whole blood was collected by terminal bleeds from the orbital sinus into K2EDTA microtainers for plasma preparation. Lung tissue (50-100 mg) were extracted with T-PER tissue protein extraction reagent (Thermo Fisher) at 1/20 (wight/ volume of reagent) with protease inhibitor. Lung tissue samples were homogenized using FastPrep-24 5G Instrument, centrifuged (10,000 × g, 5 minutes), and supernatants were analyzed. For immunoblot analysis of protein degradation, samples were prepared in reducing (for ACE2 blots) or non-reducing (for human IgG1 blots) SDS load dye and separated by polyacrylamide gel electrophoresis, transferred to PVDF membrane, blocked with 5% skim milk, and stained with 1:2000 HRP-conjugated anti-human ACE2 (clone OTI1D2, Origene) or 1:5000 HRP-conjugated donkey anti-human IgG (Jackson Immuno Research) in Tris-buffered saline containing 0.1% Tween 20 and 1% skim milk. Blots were developed with Clarity ECL Substrate (Bio-Rad) and imaged on a ChemiDoc XRS+ system (Bio-Rad).

### *In vitro* binding assays to S variants

Expi293F cells were transfected with pCEP4-myc-S plasmids as described above. 4 volumes of culture were centrifuged (600 × g, 4 °C, 120 s), the pellet was washed with 0.8 volumes ice-cold PBS-BSA, and then the cells were resuspended in 1 volume PBS-BSA. Serial dilutions of purified sACE2 or mAb proteins were prepared in round-bottomed 96-well trays using PBS-BSA as the diluent. An equal volume of cells and protein solution were then mixed and incubated at 4 °C on a rotator for 30 minutes. Plates were centrifuged (600 × g, 4 °C, 90 s) and supernatant removed. Cells were washed twice with ice-cold PBS-BSA, then resuspended in PBS-BSA containing 1:100 anti-HIS-FITC (chicken polyclonal, Immunology Consultants Laboratory) plus 1:250 anti-myc Alexa 647 (clone 9B11, Cell Signaling Technology) for detection of bound sACE2-8h proteins, or 1:250 anti-human IgG-APC (clone HP6017, BioLegend) plus 1:150 anti-MYC-FITC (chicken polyclonal, Immunology Consultants Laboratory) for detection of bound sACE2_2_-IgG1 or mAbs. Plates were incubated another 30 minutes on a rotator at 4 °C, centrifuged, cell pellets were washed twice, and finally cells were resuspended in PBS-BSA for analysis on a BD Accuri C6 flow cytometer. Data were analyzed using instrument software. In all analyses, cells were gated by side scatter-forward scatter to remove debris. To assess protein binding, cells positive for the myc tag on S were gated and the mean fluorescence signal for bound sACE2 within the myc-positive gate was measured (**Figure S6A, C-D**). To assess surface levels of S, the mean fluorescence signal for the myc tag was measured for the entire cell population. Background fluorescence of negative controls was subtracted to calculate Δ mean fluorescence. All replicates were from independent, distinct samples.

### MD simulations

For apo and RBD-bound complexes, protein structures were obtained from PDB 6M17 ^70^. Extracellular ACE2 residues 21-730 were considered in the simulations. For apo simulations, the RBD was removed from the system. Mutations were introduced using PyMOL (https://pymol.org/2/) and the systems were solvated using TIP3P water and 150 mM NaCl using PACKMOL ^71^. Protein residues and glycosylation sites were parameterized using AMBER ff14SB ^72^ and GLYCAM06 ^73^ force fields. The parameters for the Zn^2+^ center of ACE2 were collected from cationic dummy atom (CADA) method ^74^.

The systems were first minimized and equilibrated before production runs. Minimization was performed with steepest descent and conjugate gradient algorithm for 15000 steps. The minimized systems were heated in NVT ensemble with backbone atoms restrained with a restraining force (k(x)^2^) with a spring constant 10 kcal/(mol-Å^2^). The temperature was increased to 300 K in two steps (0 K to 10 K, 10 K to 300 K). Each step was performed for 2 ns. To increase the pressure of the system to 1 bar, the heated systems were simulated in NPT ensemble keeping the restraint on the backbone for another 2 ns. After this step, restraints were removed to equilibrate the system at 300 K and 1 bar for 44 ns. To maintain the temperature of the systems, Langevian dynamics were used with 2 ps^-1^ collision frequency. Berendsen barostat was used to maintain the pressure. 2 fs timestep is used for the simulation. To avoid instability of the bonds involving hydrogen atoms, SHAKE algorithm ^75^ was used. To consider non-bonded interactions, a 10 Å cutoff distance was selected. Long-range electrostatic interactions were considered by the Particle Mesh Ewald method. Minimization and equilibration were done using the AMBER18 engine.

After equilibration, production runs were performed with adaptive sampling to utilize parallel computing power. Adaptive sampling ^76^ is a parallel iterative process to sample the protein conformational space by running many short length simulations. In adaptive sampling, protein conformational space is characterized with important features like distances, angles, and root mean square deviation (RMSD), and the feature space is clustered into microstates. The starting points for the next round of simulation are selected from the least populated states. The first 5 rounds of adaptive sampling were performed using the AMBER engine on Blue Waters supercomputing facility. The features selected for the adaptive sampling were backbone RMSD. Data collected over 5 rounds of adaptive sampling were clustered into 500 clusters, from which 500 structures were selected to submit to Folding@Home ^77^. Each structure was given 20 different starting velocities to begin 20 different trajectories. Simulations were run in OPENMM^78^ engine. For each apo system, ∼ 400 µs of data were collected, and for each RBD-bound complex system, ∼530 µs of data were gathered.

A Markov state model is built on MD data to get the unbiased estimates of the canonical ensemble of interface movements. The MSM calculates the canonical ensemble distribution from the eigenvector space of the transition probability matrix. To estimate the transition probability matrix, we featurized the MD simulation data with inter-residue distances between the human and viral protein interfaces. The featured space was linearly transformed into TICs. TIC space was clustered into microstates using k-means algorithm. The probability of a jump between one microstate (i) to another (j) with a particular lag time (τ) corresponds to the T_ij_ element of the transition probability matrix, **T**. Feature calculations were performed with MDtraj. TIC calculation, clustering, and MSM building steps were done using PyEMMA ^79^. Lag time for MSM was selected by logarithmic convergence of implied timescales. To build the optimal MSM, VAMP2 scores of MSMs were calculated using different hyperparameters (TIC variance and numbers of clusters). Feature calculations and analysis of the trajectories were performed using MDtraj ^80^ and CPPtraj ^81^. For trajectory visualization, VMD software was used ^82^.

### Anti-SARS-CoV-2 Monoclonal Antibodies

Sequences for monoclonal antibodies that had received Emergency Use Authorization from the U.S. Food and Drug Administration were pulled from the KEGG database (Accession No. REGN10933, D11938; REGN10987, D11939; VIR-7831, D12014; LY-CoV555, D11936). H and L chains were cloned with CD5 leader sequences into pcDNA3.1(+) and expressed in Expi293F cells. Proteins were purified using KancapA resin (Kaneka Corporation) and size exclusion chromatography with PBS as the running buffer. Peak fractions at the expected MW of ∼150 kD were concentrated, frozen in liquid nitrogen, and stored at -80 °C. Concentrations were determined using absorbance at 280 nm and the calculated extinction coefficient for mature protein.

### Statistics

Quantification of replicate experiments is presented as the mean ± SEM. The student t-test, one-way and two-way ANOVA with Bonferroni post-tests were used to determine statistical significance, with a P value threshold of less than 0.05. Significance levels are indicated in the figures as *P < 0.05, **P < 0.01, and ***P < 0.001. Based on our experience, we expect changes in the gene/protein expression and function measurements to be detected with 3 mice per group, so the effect size was determined as n = 3 or n > 3. The variance between the groups that are being statistically compared was similar.

### Study Approval

All aspects of this study were approved by the office of Environmental Health and Safety at University of Illinois at Chicago prior to the initiation of this study. Working with SARS-CoV-2 was performed in a BSL-3 laboratory by personnel equipped with powered air purifying respirators.

### Data Availability

MD data with features for MSM building and an embedded Box link are deposited on GitHub (https://github.com/ShuklaGroup/ACE-RBD_simulation_data_script.git). Other data are available from the corresponding authors on reasonable request.

## AUTHOR CONTRIBUTIONS

L.Z., J.R., E.P, and A.B.M conceived of the project and designed the experiments; L.Z. performed experiments involving the SARS-CoV-2 isolate WA-1/2020 (U.S.A.) and the P.1. variant; L.M.C. and L.R. generated the protocols to propagate and evaluate the SARS-CoV-2 variants; S.X. performed the pseudovirus experiments; M.C. and D.S. analyzed TL mutation predictions; S.D. and D.S performed MD simulation; K.K.C. performed protein purifications, western blots of serum/lung samples, ACE2 ELISAs, ACE2 catalytic assay, and cloning; T.F., K.L.B. and M.L. performed pharmacokinetic studies; A.F.G. analyzed genotyping qPCR and Western blot data; L.Z., E.P, and J.R. wrote the initial draft of the manuscript. All authors reviewed the manuscript and provided feedback. L.Z., E.P, J.R. and A.B.M. wrote the final manuscript with editing and review.

## ACKNOWLEDGMENTS

This work was supported in part by NIH grants P01-HL60678, R01-HL154538, R01-HL149300, R01-HL118068, R01-HL157489 and R01-HL152515 to A.B.M and J.R.; R43-AI162329 to E.P. and K.K.C.; R01HL157489 to L.Z.; T32-HL007829 to A.B.M; support to D.S. from the C3.ai Digital Transformation Institute Research Award provided by C3.ai Inc. and Microsoft Corporation; computational time from Folding@home; as well as by intramural funds of the University of Illinois College of Medicine. The following reagents were obtained through BEI Resources, NIAID, NIH: 2019n-CoV/USA_WA1/2019 isolate of SARS-CoV-2 (NR-52281) and Isolate hCoV-19/Japan/TY7-503/2021 (P.1), NR-54982. We thank Dr. Maria Swerdlov at the Histology Tissue Core for the support and Dr. Justin M. Richner at Department of Microbiology and Immunology for providing BSL3 training. We thank Dr. David Ho (Columbia University) for providing the SARS-CoV-2 pseudovirus.

