## Supplemental Figures S1-S6 and Tables S1-S3 for "Engineered High-Affinity ACE2 Peptide Mitigates ARDS and Death Induced by Multiple SARS-CoV-2 Variants"

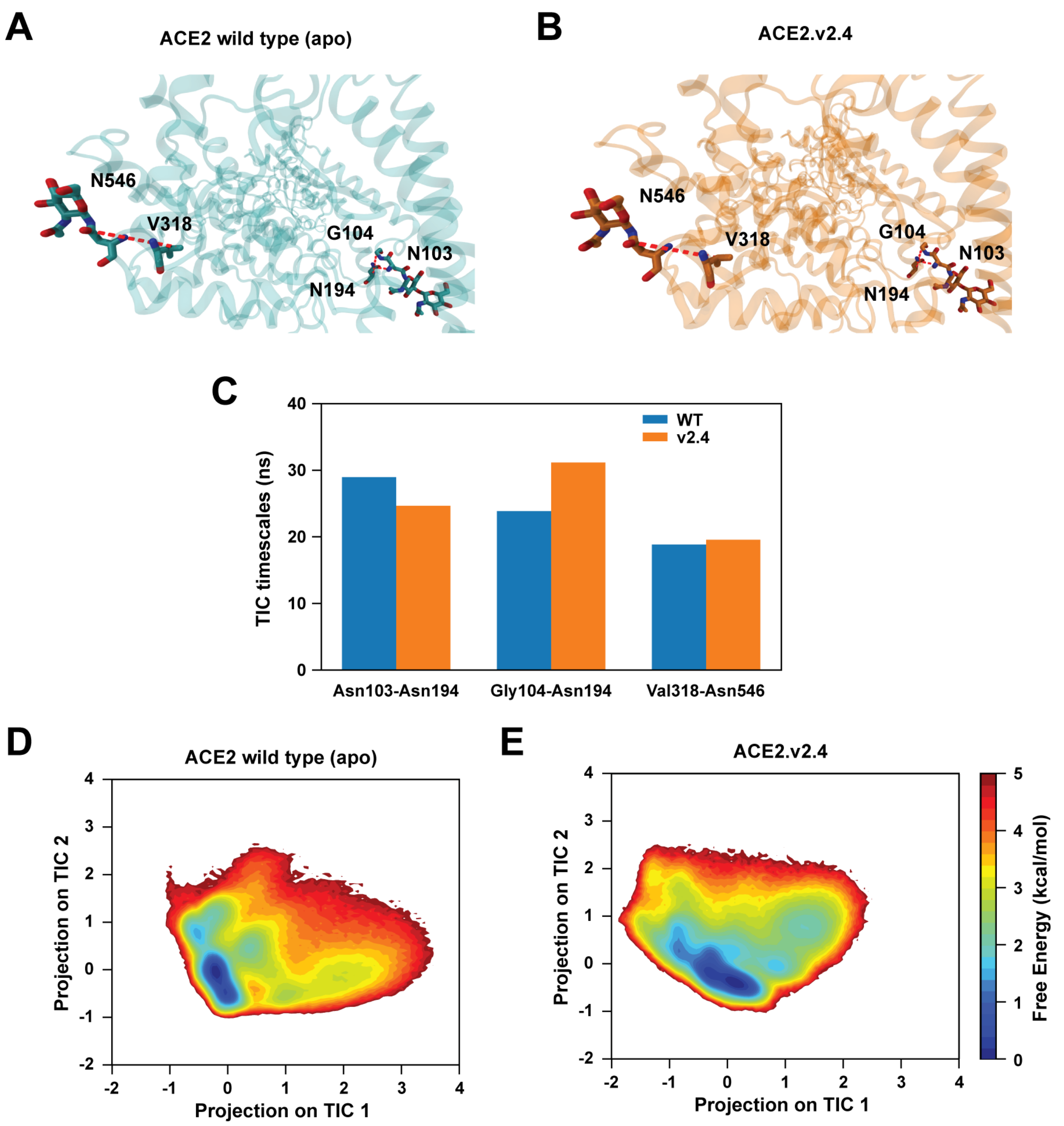


**Figure S1. Surface dynamics of unbound wild type ACE2 and ACE2.v2.4 are similar.** (**A** & **B**) Based on simulations of apo ACE2, the three distance features are shown that have highest correlation to the slowest TICs for wild type (**A**) and v2.4 (**B**) proteins. Proteins are shown as blue and orange cartoons, respectively. Important residues are represented as sticks and measured distances are shown as broken red lines. (**C**) TIC timescales for the three slowest TIC components. WT, blue; v2.4, orange. (**D** & **E**) Free energy landscapes for the projections of TIC 1 and TIC 2 for wild type (**D**) and v2.4 ACE2 (**E**).

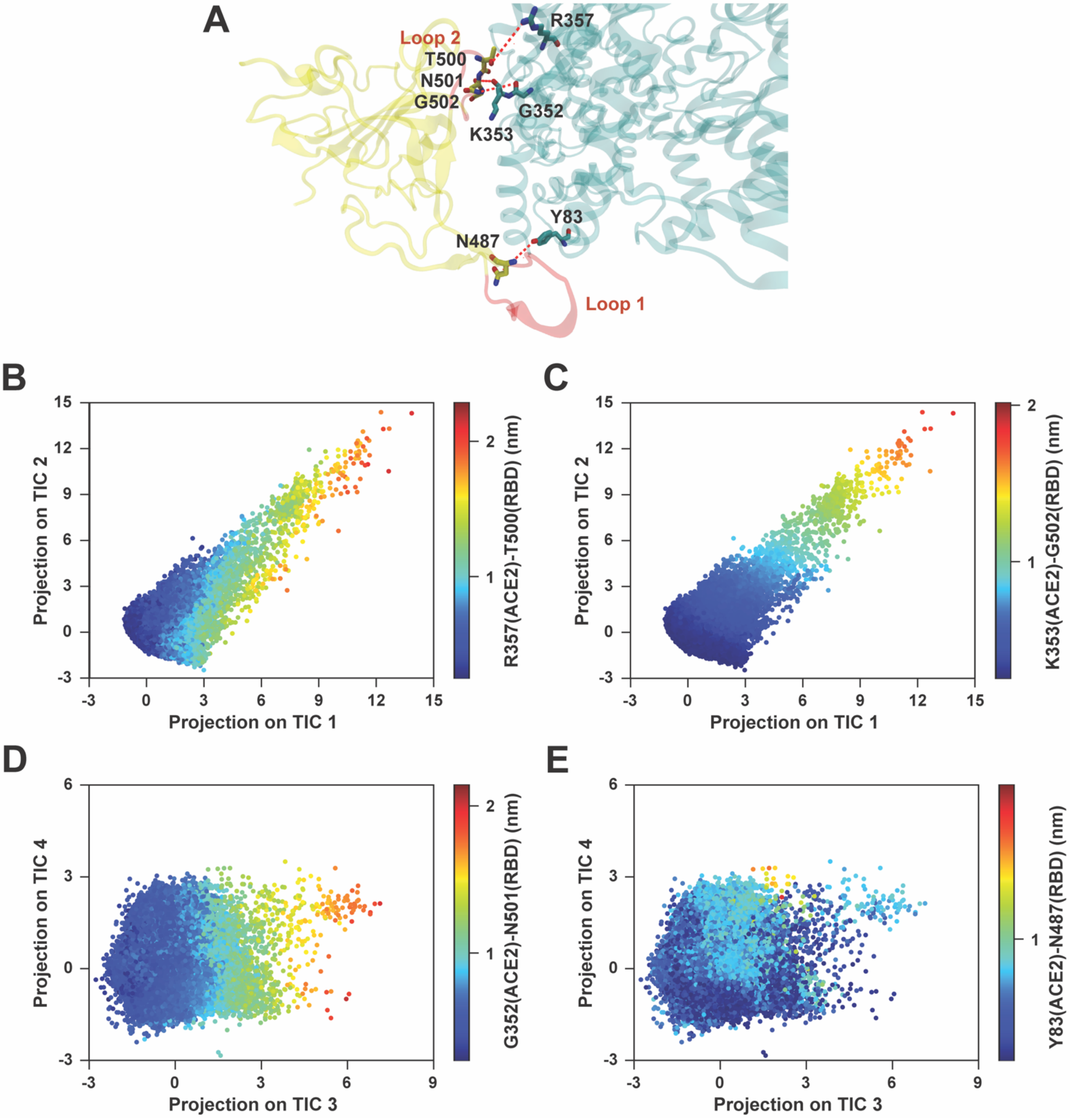


**Figure S2. TIC correlations with interface residue distances.** (**A**) For the complex of wild type ACE2 (blue) with RBD (yellow), distance features correlated with the slowest TICs are represented. RBD loops 1 and 2 are red. (**B** & **C**) Projection of TIC 1 coordinate with respect to TIC 2. Color scales are defined based on interaction distances with highest correlation to TIC 1 (**B**) or TIC 2 (**C**). (**D** & **E**) Projection of TIC 3 coordinate with respect to TIC 4. Color scales are defined based on interaction distances with highest correlation to TIC 3 (**D**) or TIC 4 (**E**).

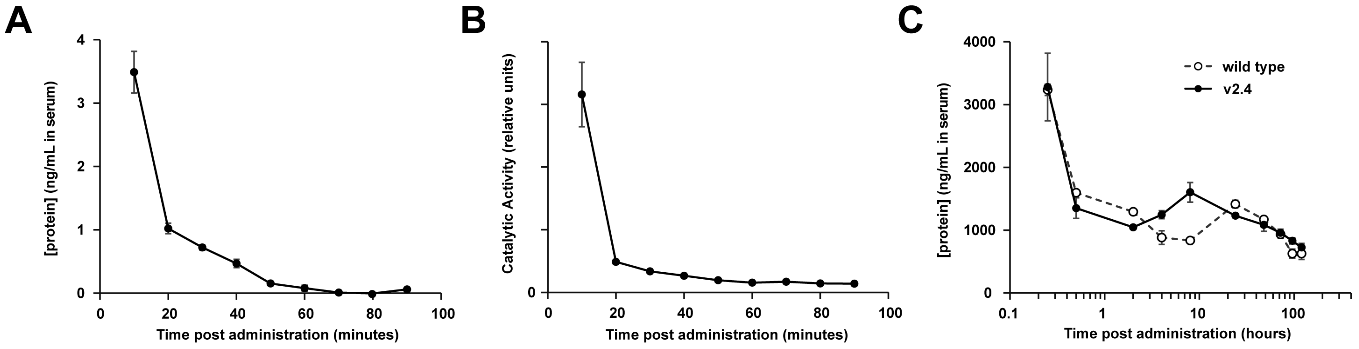


**Figure S3. Serum PK of IV administered sACE2$_2$.v2.4 and IgG1-fusion proteins.** (**A** & **B**) Unfused sACE2$_2$.v2.4 was injected in the tail veins of mice (3 male and 3 female per time point; 0.5 mg/kg). Serum was analyzed (**A**) by ACE2 ELISA and (**B**) for proteolytic activity towards a fluorogenic substrate. (**C**) IV administration of 2.0 mg/kg wild type sACE2$_2$-IgG1 (white circles) or sACE2$_2$.v2.4-IgG1 (black circles) in 3 male mice per time point. Protein in serum was quantified by human IgG1 ELISA. Data are mean ± SEM.

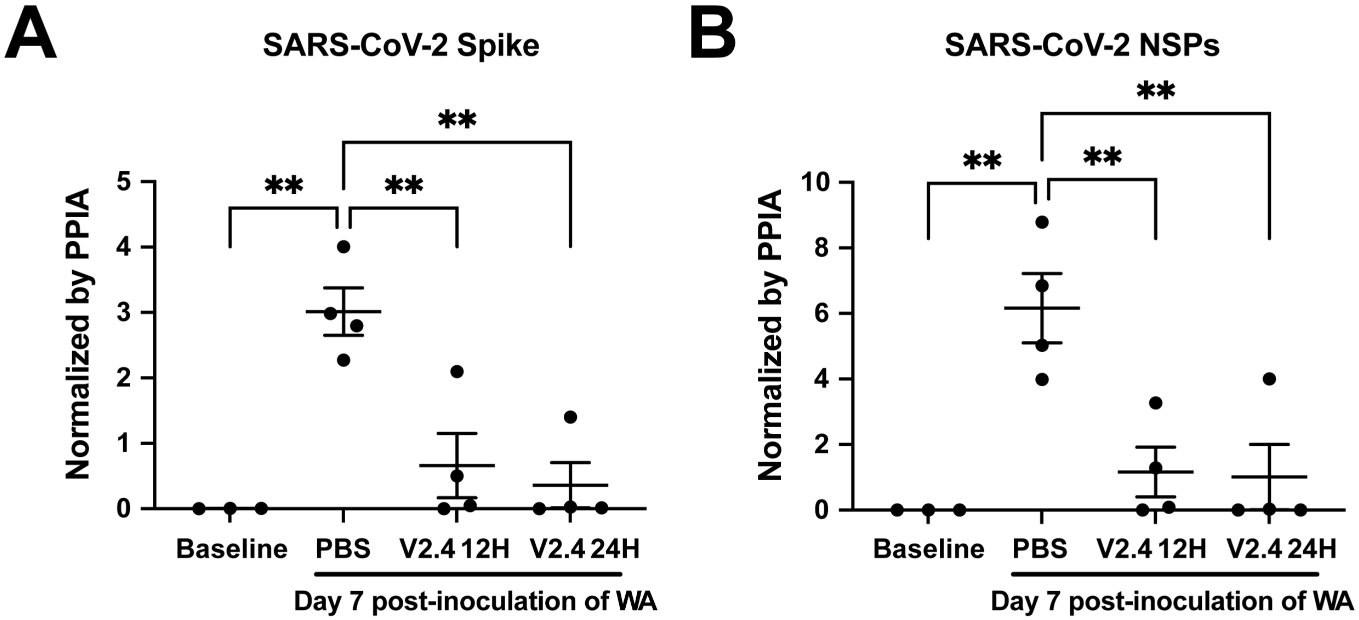


**Figure S4.** (**A** & **B**) Viral load of the lungs harvested at Day 7 post-inoculation with SARS-CoV-2 strain WA-1/2020 as measured by real-time quantitative PCR for the mRNA expression level of SARS-CoV-2 Spike and SARS-CoV-2 NSPs. N=4
Data are presented as mean ± SEM. **: P<0.01 by one-way ANOVA.

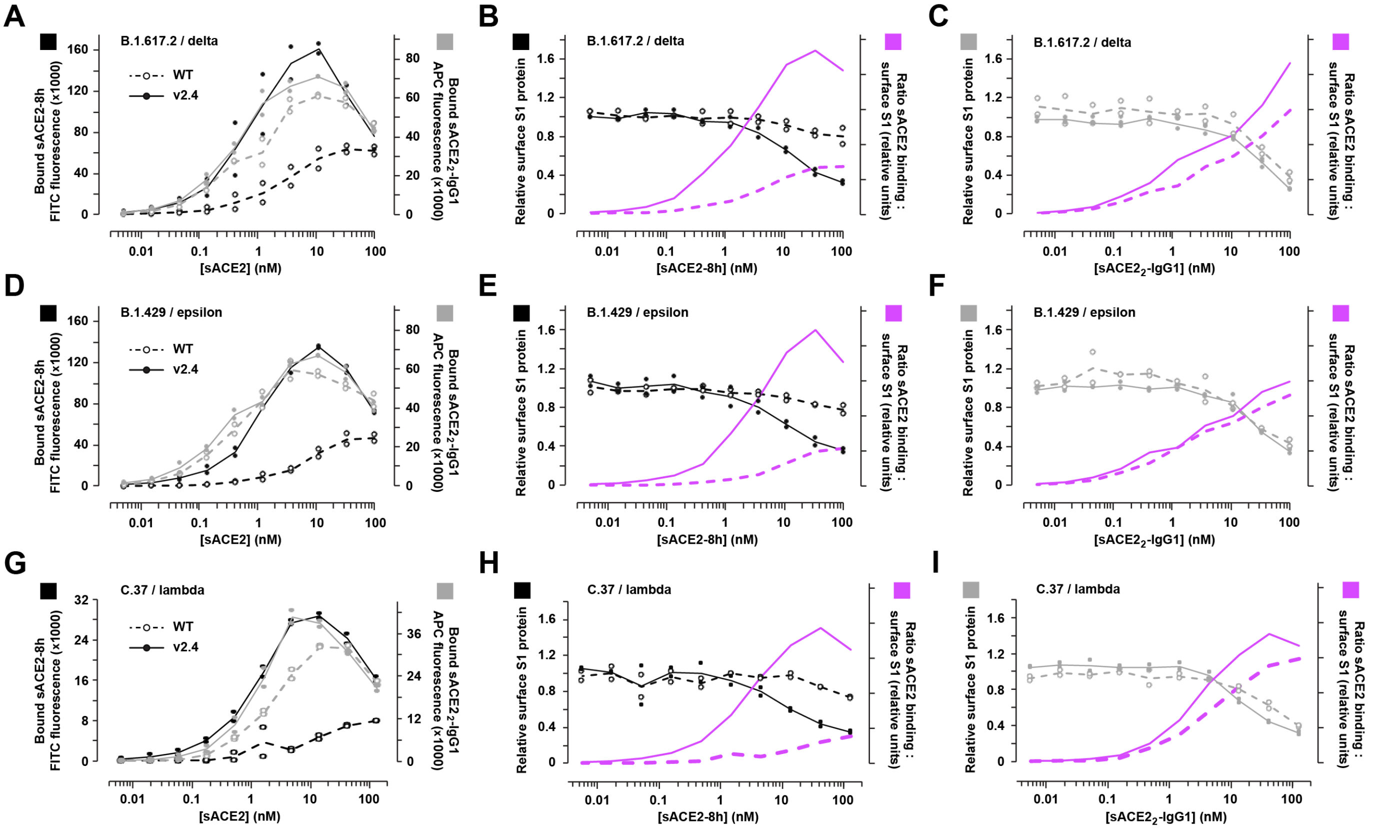

**Figure S5. Decoy receptors diminish S1 at a membrane surface.**
Human Expi293F cells expressing myc-S from the B.1.617.2 (**A-C**), B.1.427/B.1.429 (**D-F**), or C.37 (**G-I**) lineages were incubated with monomeric sACE2-8h (black) or dimeric sACE2$_2$-IgG1 (grey), washed, and then stained for flow cytometry (N = 2). (**A, D, G**) Bound wild type (dashed lines, open circles) and v2.4 (solid lines, filled circles) sACE2 proteins. (**B-C, E-F, H-I**) Relative surface S1 protein based on detection of the myc tag, after incubating cells with sACE2-8h (**B, E, H**) or sACE2$_2$-IgG1 (**C, F, I**). The ratio of bound decoy receptor to surface S1 is shown in magenta.

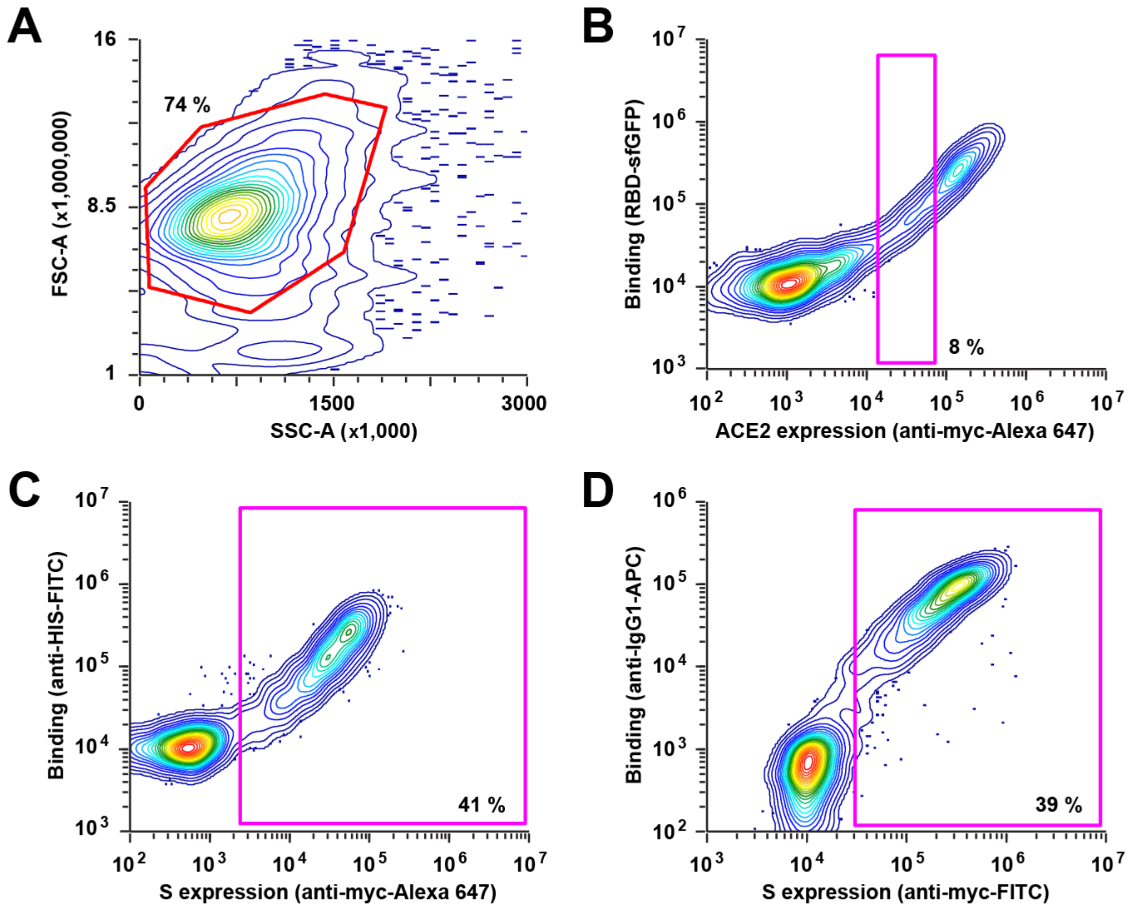


**Figure S6. Gating strategies for measuring ACE2-S binding by flow cytometry.**
(**A**) Human Expi293F cells transfected with myc-tagged S or ACE2 proteins were washed, incubated with soluble ACE2 and mAbs (to measure binding to transmembrane S) or soluble RBD-sfGFP (to measure binding to transmembrane ACE2), and stained with fluorescent antibodies to detect surface S or ACE2 expression and bound soluble proteins. Cells were analyzed by flow cytometry and in all cases the main population was gated (red) first by forward scatter (FSC) and side scatter (SSC). (**B**) To measure binding of RBD-sfGFP to myc-ACE2 expressing mutants, mean sfGFP fluorescence was measured in cells gated (magenta) for a consistent level of ACE2 expression. This controls for differences in surface expression levels between ACE2 mutants. (**C**) To measure binding of monomeric sACE2-8h proteins to myc-S expressing cells, mean fluorescence for bound protein is reported within the myc-positive gate (magenta). Surface S expression was measured by recording the mean anti-myc-Alexa 647 fluorescence signal for the entire population. (**D**) To measure binding of dimeric $sACE2_2$-IgG1 or anti-SARS-CoV-2 mAbs to myc-S expressing cells, mean fluorescence for bound protein is reported for the myc-positive gate (magenta). Surface S expression is based on mean fluorescence for the anti-myc-FITC signal in the entire population.
For all panels, representative samples are shown with percentages of cells in the gates indicated.

| Table S1. Prophylaxis studies of monoclonal antibodies and ACE2 decoys. | | | | | | | | |
|---|---|---|---|---|---|---|---|---|
| **Reference (DOI)** | **Manuscript Title** | **Administered Proteins** | **Animal Model** | **SAR-CoV2 Variant** | **Dose of Virus** | **Administration of Test Article [1]** | **Assessment** | **Conclusions** |
| This Study | Engineered High-Affinity ACE2 Peptide Mitigates ARDS and Death Induced by Multiple SARS-CoV-2 Variants | Engineered ACE2 decoy: sACE22.v2.4-IgG1 | K18-hACE2 transgenic mice | USA-WA1/2020 | 10^4 PFU intranasally | 10 mg/kg i.v. 12 h prior to infection | Daily weight loss, daily survival, lung vascular leakage and edema at d 7 | Animals were fully protected. |
| 10.1101/2020.06.15.152157 | Novel ACE2-IgG1 fusions with improved in vitro and in vivo activity against SARS-CoV2 | Wild type or catalytically inactive sACE2-IgG1(LALA) | AdV-hACE2 transduced mice | Unspecified. Virus isolate is from early 2020. | 5 × 10^4 to 2 × 10^5 PFU intranasally | 15 mg/kg i.v. 4 h prior to infection | Viral load in lungs on d 3 | Wild type sACE2-IgG1 had no effect. Catalytically inactive sACE2-IgG1 with slightly increased SARS-CoV-2 affinity caused moderate decrease in virus load. |
| 10.1371/journal.ppat.1009544 | Intranasal gene therapy to prevent infection by SARS-CoV-2 variants | Engineered ACE2 decoy: CDY14HL-Fc4 | K18-hACE2 transgenic mice | USA-WA1/2020 | 2.8 × 10^2 PFU intranasally | Gene therapy: i.n. delivery of AAV expressing CDY14HL-Fc4 7 days prior to infection | Daily weight loss, viral RNA and histopathology on days 4 and 7 | Small to moderate reductions in weight loss, viral RNA levels, and lung pathology. |
| 10.1038/s41586-021-03720-y | In vivo monoclonal antibody efficacy against SARS-CoV-2 variant strains | mAbs: 2B04+47D11, S309+S2E12, COV2-2130+COV2-2196, REGN10933+REGN10987, or LY-CoV555 | K18-hACE2 transgenic mice | WA1/2020 N501Y/D614G, B.1.1.7, Wash-B.1.351 and Wash-B.1.1.28 | 10^3 FFU intranasally | 0.2 mg/kg and 2 mg/kg i.p. 24 h prior to infection | Daily weight loss, viral loads in tissues at day 3 or 6, cytokine analysis | Limited, variant-specific protection by 2B04+47D11 and LY-CoV555. Reductions in viral RNA were diminished for B.1.351 virus, especially for REGN10933+REGN10987 and COV2-2130+COV2-2196. Overall, cocktails were protective. |
| 10.1038/s41586-021-03720-y | In vivo monoclonal antibody efficacy against SARS-CoV-2 variant strains | mAbs: COV2-2130+COV2-2196 | Syrian hamster | Wash-B.1.351 and WA1/2020 D614G | 5 × 10^5 FFU intranasally | 4 mg/kg and 10 mg/kg i.p. 24 h prior to infection | Daily weight loss, viral loads in tissues at day 4, cytokine analysis | Animals were protected. Viral RNA remained high in nasal washes. |
| 10.1084/jem.20201993 | Antibody potency, effector function and combinations in protection from SARS-CoV-2 infection in vivo | mAbs: C002, C104, C105, C110, C119, C121-LS, C135-LS, and C144-LS | Wild type mice | SARS-CoV-2 MA | 10^5 PFU intranasally | 8mg/kg i.p. 12 h prior to infection of individual mAbs, and 16 / 5.3 / 1.8 mg/kg i.p. for cocktails (C135-LS/C121-LS and C135-LS/C144-LS) | Viral load in lungs at day 2. This is a non-lethal infection model. | Prophylactic efficacy in vivo did not correlate well with in vitro neutralization and required effector functions. Cocktail can out-perform monotherapy at a lower dose (5.3 mg/kg). Protection was partial at the lowest dose of 1.8 mg/kg. |
| 10.1084/jem.20201993 | Antibody potency, effector function and combinations in protection from SARS-CoV-2 infection in vivo | mAbs: C135-LS+C144-LS | Syrian hamster | Unspecified | 2.6 × 10^4 PFU intranasally | 20 / 6 / 2 mg/kg i.p. 24 h prior to infection | Viral load in lungs at d 3. | All doses substantially reduced viral load at d 3. |
| 10.1038/s41586-020-2548-6 | Potently neutralizing and protective human antibodies against SARS-CoV-2 | mAbs: COV2-2196, COV2-2130, and COV2-2196+COV2-2130 | AdV-hACE2 transduced mice | USA_WA1/2020 | 4 × 10^5 PFU intranasally | 10 mg/kg i.p. 24 prior to infection | Daily weight loss, histopathology at day 7, viral RNA at day 7, inflammatory cytokines at day 7 | Partial protection from weight loss and infection. |
| 10.1038/s41586-020-2548-6 | Potently neutralizing and protective human antibodies against SARS-CoV-2 | mAbs: COV2-2196, COV2-2130, and COV2-2196+COV2-2130 | Wild type mice | SARS-CoV-2 MA | 10^5 PFU intranasally | 10 mg/kg i.p. 6 h prior to infection | Viral RNA in lungs at d 2 | Reduction in virus replication |
| 10.1038/s41586-020-2548-6 | Potently neutralizing and protective human antibodies against SARS-CoV-2 | mAbs: COV2-2196 or COV2-2130 | Rhesus macaque | USA_WA1/2020 | 1.1 × 10^4 PFU intranasally and intratracheally | 50 mg/kg i.v. 3 days prior to infection | Viral RNA in bronchoalveolar lavage and nasal swabs | Reduction in virus replication |
| 10.1038/s41586-020-2381-y | A human neutralizing antibody targets the receptor-binding site of SARS-CoV-2 | mAb: CB6 | Rhesus macaques | BetaCoV/Wuhan/IVDC-HB-envF13/2020 | 10^5 TCID50 intratracheally | 50 mg/kg i.v. 24 h prior to infection | Viral RNA in throat swabs, histopathology at day 5 | Blocked virus replication |
| 10.1101/2021.03.09.434607 | The dual function monoclonal antibodies VIR-7831 and VIR-7832 demonstrate potent in vitro and in vivo activity against SARS-CoV-2 | mAb: VIR-7831 | Syrian hamster | USA-WA1/2020 | 7.4 × 10^4 TCID50 intranasally | 30 / 5 / 0.5 / 0.05 mg/kg 24 h or 15 / 5 / 0.5 / 0.05 mg/kg 48 h i.p. prior to infection | Lung viral load at day 4, daily body weight change | VIR-7831 decreased viral load and ameliorated weight loss. Efficacy dropped substantially at doses less than 5 mg/kg. |
| 10.4049/jimmunol.2000583 | A Potently Neutralizing Antibody Protects Mice against SARS-CoV-2 Infection | Murine mAb: 2B04 | AdV-hACE2 transduced mice | USA-WA1/2020 | 4 × 10^5 FFU intranasally | 10 mg/kg 2B04 i.p. 24 h prior to infection | Daily weight, histopathology and viral load on day 4 | Ameliorated weight loss and reduced viral load |
| 10.1126/scitranslmed.abf1906 | The neutralizing antibody, LY-CoV555, protects against SARS-CoV-2 infection in nonhuman primates | mAb: LY-CoV555 | Rhesus macaque | USA-WA1/2020 | 1.1 × 10^5 PFU intranasally and intratracheally | 1 / 2.5 / 15 / 50 mg/kg i.v. 24 h prior to infection | Viral RNA in nasal and throat swabs and in lungs | All doses reduced viral RNA, especially at 2.5 mg/kg and higher. |
| 10.1016/j.cell.2020.05.025 | Potent Neutralizing Antibodies against SARS-CoV-2 Identified by High-Throughput Single-Cell Sequencing of Convalescent Patients' B Cells | mAb: BD-368-2 | hACE2 transgenic mice | Wuhan/AMMS01/2020 | 10^5 TCID50 intranasally | 20 mg/kg i.p. 24 h prior to infection | Daily weight loss, viral RNA in lungs on day 5 | Ameliorated weight loss and blocked virus replication |
| 10.1126/science.abe2402 | REGN-COV2 antibodies prevent and treat SARS-CoV-2 infection in rhesus macaques and hamsters | mAb cocktail: REGN10933+REGN10987 | Rhesus macaque | USA-WA1/2020 | 10^5 or 1.05 × 10^6 PFU intranasally and intratracheally | 0.3 / 50 mg/kg i.v. 72 h prior to infection | Viral RNA in nasal and throat swabs and BAL over 8 days, histopathology | 50 mg/kg dose was effective, 0.3 mg/kg was not. |
| 10.1126/science.abe2402 | REGN-COV2 antibodies prevent and treat SARS-CoV-2 infection in rhesus macaques and hamsters | mAb cocktail: REGN10933+REGN10987 | Syrian hamster | USA-WA1/2020 | 2.3 × 10^4 PFU intranasally | 50 / 5 / 0.5 mg/kg i.v. 48 h prior to infection | Daily weight loss, viral RNA in lungs and histopathology on day 7 | All doses prevented weight loss, reduced viral load and reduced severity of lung damage |
| 10.1126/science.abf4830 | Broad and potent activity against SARS-like viruses by an engineered human monoclonal antibody | mAb: ADG-2 | Wild type mice | SARS-CoV-MA15 or SARS2-CoV-2-MA10 | 10^3 PFU intranasally | 10 mg/kg i.p. 12 h prior to infection | Daily weight loss, lung function and injury, lung viral load at days 2 and 4 | Prophylaxis completely protected mice |

[1] In cases where mice were administered a defined mass of test article, mouse weight is approximated to be 20 g for calculating a dose in mg/kg.

| Table S2. Therapeutic studies of monoclonal antibodies and ACE2 decoys. | | | | | | | | |
|---|---|---|---|---|---|---|---|---|
| Reference (DOI) | Manuscript Title | Administered Proteins | Animal Model | SAR-CoV2 Variant | Dose of Virus | Administration of Test Article [1] | Assessment | Conclusions |
| This Study | Engineered High-Affinity ACE2 Peptide Mitigates ARDS and Death Induced by Multiple SARS-CoV-2 Variants | Engineered ACE2 decoy: sACE22.v2.4-IgG1 | K18-hACE2 transgenic mice | USA-WA1/2020 and Japan/TY7-503/2021 (P.1 variant) | 10^4 PFU intranasally | 10 mg/kg i.v. daily for 7 days beginning 12 h post-infection, or 15 mg/kg i.v. daily for 7 days beginning 24 h post infection, | Daily weight loss, daily survival, lung vascular leakage, edema and histopathology at days 6 or 7 and 14, viral load in lungs on day 6 or 7 | Animals were partially protected from weight loss, death, vascular leakage, and lung pathology. Viral burden substantially reduced. Animals fully recovered in 14 days. P.1 virus had a faster disease course and required treatment to begin at 12 h for survival, although later treatment still reduced lung vascular leakage, edema, and viral burden. |
| 10.1038/s41467-021-24013-y | Engineered ACE2 receptor therapy overcomes mutational escape of SARS-CoV-2 | Engineered ACE2 decoy: 3N39v2-Fc | Syrian hamster | Japan/TY/WK-521/2020 | 10^6 PFU intranasally | 20 mg/kg i.p. 2 h post infection, smaller study where administered 48 hours post infection | Daily body weight, on day 5 assessment of viral load, inflammatory gene expression, and histopathology | Treated (2 h post-infection) animals recovered in body weight by day 5 and had reduced inflammatory gene expression and viral load. Treatment 2 days post-infection gave smaller decreases in viral load and inflammatory gene expression, no data provided on weight loss. |
| 10.1038/s41586-021-03720-y | In vivo monoclonal antibody efficacy against SARS-CoV-2 variant strains | mAbs: 2B04+47D11, S309+S2E12, COV2-2130+COV2-2196, REGN10933+REGN10987, LY-CoV555+LY-CoV016, or LY-CoV555 | K18-hACE2 transgenic mice | WA1/2020 N501Y/D614G and Wash-B.1.351 | 10^3 FFU intranasally | 10 mg/kg i.p. 24 h post infection | Daily weight loss, viral loads in tissues at day 6, histopathology | Except for treatment with LY-CoV555 or LY-CoV555+LY-CoV016 in B.1.351 infected mice, animals were protected. Unlike prophylaxis, in vitro neutralization efficacy did not correlate well with therapeutic efficacy, indicating the importance of other effector functions. |
| 10.1084/jem.20201993 | Antibody potency, effector function and combinations in protection from SARS-CoV-2 infection in vivo | mAbs: C135-LS+C144-LS | Syrian hamster | Unspecified | 2.6 × 10^4 PFU intranasally | 40 / 12 / 4 mg/kg i.p. 12 h post infection | Viral load in lungs at d 3. | All doses substantially reduced viral load at d 3. |
| 10.1038/s41586-020-2548-6 | Potently neutralizing and protective human antibodies against SARS-CoV-2 | mAbs: COV2-2196+COV2-2130 | AdV-hACE2 transduced mice | USA_WA1/2020 | 4 × 10^5 PFU intranasally | 20 mg/kg i.p. 12 h post infection | Viral RNA in lungs at day 2, inflammatory cytokines | Reduction in virus replication and inflammatory cytokines |
| 10.1038/s41586-020-2548-6 | Potently neutralizing and protective human antibodies against SARS-CoV-2 | mAbs: COV2-2196, COV2-2130, and COV2-2196+COV2-2130 | Wild type mice | SARS-CoV-2 MA | 10^5 PFU intranasally | 20 mg/kg i.p. 12 h post infection | Viral RNA in lungs at d 2 | Reduction in virus replication |
| 10.1038/s41467-020-20602-5 | A therapeutic neutralizing antibody targeting receptor binding domain of SARS-CoV-2 spike protein | mAb: CT-P59 | Ferret | NMC-nCoV02 | 10^5.5 - 10^5.8 TCID50 intranasally and intratracheally | 3 / 30 mg/kg i.v. 24 h post infection | Viral load in nasal washes, histopathology at days 3 and 7 | 30 mg/kg dose reduced viral load. Lower dose much less efficacious. |
| 10.1038/s41467-020-20602-5 | A therapeutic neutralizing antibody targeting receptor binding domain of SARS-CoV-2 spike protein | mAb: CT-P59 | Syrian hamster | NMC-nCoV02 | 6.4 × 10^4 PFU intranasally | 15 / 30 / 60 / 90 mg/kg i.p. 24 h post infection | Viral load at days 3 and 5 | Reduced virus replication. Less efficacious at lowest dose (15 mg/kg) |
| 10.1038/s41467-020-20602-5 | A therapeutic neutralizing antibody targeting receptor binding domain of SARS-CoV-2 spike protein | mAb: CT-P59 | Rhesus macaque | NMC-nCoV02 | 6.4 × 10^4 PFU intranasally | 45 / 90 mg/kg i.v. 24 h post infection | Viral load in nasal and throat swabs and in lungs | Reduced virus replication in upper respiratory tract but no change in viral RNA in lungs. |
| 10.1038/s41586-020-2381-y | A human neutralizing antibody targets the receptor-binding site of SARS-CoV-2 | mAb: CB6 | Rhesus macaques | BetaCoV/Wuhan/IVDC-HB-envF13/2020 | 10^5 TCID50 intratracheally | 50 mg/kg i.v. administered twice (days 1 and 3) post infection | Viral RNA in throat swabs, histopathology at day 5 | Reduced virus replication |
| 10.1016/j.cell.2020.05.025 | Potent Neutralizing Antibodies against SARS-CoV-2 Identified by High-Throughput Single-Cell Sequencing of Convalescent Patients' B Cells | mAb: BD-368-2 | hACE2 transgenic mice | Wuhan/AMMS01/2020 | 10^5 TCID50 intranasally | 20 mg/kg i.p. 2 h post infection | Daily weight loss, viral RNA in lungs on day 5 | Ameliorated weight loss and reduced virus replication |
| 10.1126/science.abe2402 | REGN-COV2 antibodies prevent and treat SARS-CoV-2 infection in rhesus macaques and hamsters | mAb cocktail: REGN10933+REGN10987 | Rhesus macaque | USA-WA1/2020 | 10^6 PFU intranasally and intratracheally | 25 / 150 mg/kg i.v. 24 h post infection | Viral RNA in nasal and throat swabs and BAL over 7 days, histopathology | Partially effective at reducing viral load. |
| 10.1126/science.abe2402 | REGN-COV2 antibodies prevent and treat SARS-CoV-2 infection in rhesus macaques and hamsters | mAb cocktail: REGN10933+REGN10987 | Syrian hamster | USA-WA1/2020 | 2.3 × 10^4 PFU intranasally | 50 / 5 / 0.5 mg/kg i.v. 24 h post infection | Daily weight loss | 5 and 50 mg/kg doses ameliorated weight loss |
| 10.1126/science.abf4830 | Broad and potent activity against SARS-like viruses by an engineered human monoclonal antibody | mAb: ADG-2 | Wild type mice | SARS2-CoV-2-MA10 | 10^3 PFU intranasally | 10 mg/kg i.p. 12 h post infection | Daily weight loss, lung function and injury, lung viral load at days 2 and 4 | Partial amelioration of pathology of varying significance, virus levels reduced at day 4 |

[1] In cases where mice were administered a defined mass of test article, mouse weight is approximated to be 20 g for calculating a dose in mg/kg.

| Table S3. qPCR primer sequences | | |
|---|---|---|
| Target gene | Forward primer | Reverse primer |
| SARS- CoV-2 Spike | GCTGGTGCTGCAGCTTATTA | AGGGTCAAGTGCACAGTCTA |
| SARS- CoV-2 NSP | CAATGCTGCAATCGTGCTAC | GTTGCGACTACGTGATGAGG |
| Cdh5 | GTCGATGCTAACACAGGGAATG | AATACCTGGTGCGAAAACACA |
| PPIA | GGCAAATGCTGGACCAAACAC | TTCCTGGACCCAAAACGCTC |
